# Striatal ensembles specify and control ongoing actions at fine motor resolution

**DOI:** 10.64898/2025.12.03.692128

**Authors:** Ines Rodrigues-Vaz, Vivek R. Athalye, Helio F.M. Rodrigues, Darcy S. Peterka, Rui M. Costa

**Affiliations:** Zuckerman Mind Brain Behavior Institute, Departments of Neuroscience and Neurology; Columbia University; New York, USA; Champalimaud Neuroscience Programme, Champalimaud Centre for the Unknown, Lisbon, Portugal; Allen Institute, Seattle, USA; Aligning Science Across Parkinson’s (ASAP) Collaborative Research Network, Chevy Chase, USA; Kavli Institute for Brain Science, Columbia University; New York, USA

**Author notes:** Corresponding author. (RMC). These authors contributed equally to this work (Ines Rodrigues-Vaz, Vivek R. Athalye).

## Abstract

The striatum is classically thought to modulate action vigor and reinforcement, but whether it specifies and controls ongoing actions at a fine scale remains unclear. Here, we test whether striatal activity can resolve and control actions as granular as distinct patterns of muscle co-contraction within the same limb. We developed a closed-loop system combining two-photon imaging and holographic optogenetics to identify and manipulate action-specific striatal ensembles in real time. Mice performed isometric push or pull actions engaging the same forelimb muscles with different activation patterns. Ensembles of both D1- and D2-spiny projection neurons encoded action identity with equal fidelity. Causal stimulation of small ensembles biased ongoing force, but only when the ensembles encoded the ongoing action. These results show that the striatum specifies and controls ongoing actions in real time at a previously unappreciated granularity, redefining its role in motor control.

## Main Text

The brain must generate signals that enable the execution of appropriate actions, many of which demand granular motor control. Writing, for example, requires precisely coordinated muscle co-contractions in the arm to apply specific forces to a pen. Whereas motor cortices, brainstem, and spinal circuits are classically linked (*1*) to the online specification and execution of movement kinematics and force, the striatum has been primarily framed as a center for invigoration (*2–6*) and reinforcement (*5–13*) rather than as a controller of ongoing action per se. This view has been supported by the fact that Parkinson’s disease (*14–16*), where dopamine inputs to striatum are depleted, leads to deficits in action initiation (*17*), vigor (*2*), and reinforcement (*18, 19*). Alternatively, the striatum could specify actions (*20–23*), rather than merely broadcasting a more generic “go/no-go” (*24*) or vigor signal. This is consistent with observations that in disorders (*14–16*) where striatal neurons are affected, like in Huntington’s disease (*15, 25*) or dystonia (*15, 26, 27*), the execution of particular specific movements is disrupted. Recent work shows that striatal activity can encode particular whole-body (*20, 21*) and forelimb movements (*22, 23*), but it remains unclear how granular this encoding is. At one extreme, striatal activity could specify only broad action classes (e.g., body part or movement direction). At the other, it could resolve subtle differences in muscle co-contraction patterns within the same effector. Furthermore, whether these representations are epiphenomenal or causally control ongoing actions remains unknown.

Striatal projection neurons are composed of two distinct cell types, D1- and D2-spiny projection neurons (SPNs), with different dopamine receptors and projection patterns, that have been canonically assigned opponent roles in movement promotion and suppression, respectively (*15, 16, 28, 29*). However, both receive cortical inputs that carry specific movement plans (*30–34*), both are activated at movement initiation (*35, 36*), and both seem necessary for execution (*36, 37*), raising the possibility that both pathways may participate in specifying actions rather than just implementing opponent control signals.

To address these questions, we developed a real-time system to identify neuronal ensembles with a statistical model and manipulate them with cellular precision in closed-loop with behavior. Our all-optical system measured neuronal activity with two-photon calcium imaging and stimulated targeted neurons with holographic optogenetics (*38–47*) deep in the brain through a GRIN lens (*41, 46*). This approach enabled, to our knowledge, the first closed-loop identification and manipulation of action-specific ensembles in the striatum with cellular precision during ongoing actions. Further, we designed a self-paced isometric force task in which head-fixed mice executed push or pull forelimb actions on an immobile joystick, engaging both flexor and extensor muscles with different activation patterns but no overt kinematics. This task allowed us to probe whether the striatum specifies and controls ongoing actions at the granularity of distinct co-contraction patterns, and whether ensembles of D1- and D2-SPNs do so equally or with opponent roles. Together, this strategy allowed us to determine that the striatum specifies ongoing actions, with specific ensembles of both D1- and D2-SPNs controlling specific actions, as granular as distinct patterns of forelimb muscles without overt movement.

### A two-action isometric force task to study granular forelimb actions

To investigate whether the striatum encodes and controls granular actions of the same body part, we developed a task in which mice performed two forelimb actions (Fig. 1A and S4A), consisting of a push or pull force on an immobile joystick in the absence of overt movement (i.e. isometric, Methods). Mice initiated actions in a self-paced manner, in the absence of external cues. The task enabled analysis of push and pull actions with matched continuous parameters including force, duration, and licking (fig. S3A-H, Methods).

**Fig. 1.**
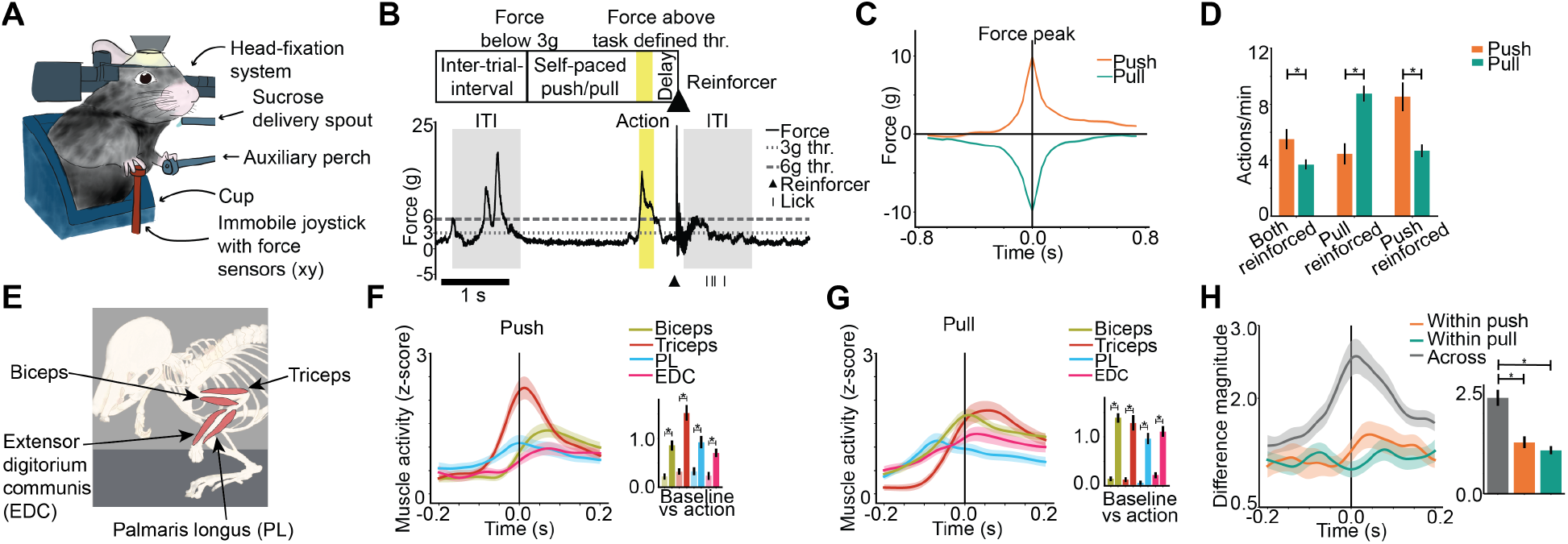
Mice perform a two-action isometric force task activating both flexor and extensor forelimb muscles with different patterns. (**A**) Task schematic. (**B**) Task events for reinforcement of self-paced actions. Trace of force applied to the joystick on an example trial. (**C**) Trial-averaged force trace of 6g cross actions time locked to force peak. By our convention, force in the push direction is positive and pull direction is negative. Data are mean+s.e.m. across 109 sessions from 8 mice. (**D**) Rate of 6g cross actions. Mice performed both actions and increased the action which led to sucrose reinforcement. Data are mean+s.e.m. from 8 mice, averaging over sessions in each block. Mice performed push more than pull when both actions were reinforced (table S1, ref. 1.1). Mice performed pull more than push when pull was reinforced (table S1, ref. 1.2). Mice performed push more than pull when push was reinforced (table S1, ref. 1.3). (**E**) Schematic of flexor and extensor forelimb muscles simultaneously recorded with EMG. (**F**) Trial-averaged z-scored EMG of 6g cross push actions locked to 6g cross, force-matched with pull actions. Each muscle significantly activated at 6g cross relative to 1 second before (table S1, ref. 1.4-1.7). (**G**) Same as (F) but for pull action, force-matched with push actions. Each muscle significantly activated at 6g cross relative to 1 second before (table S1, ref. 1.8-1.11). (**H**) Vector magnitude of EMG difference at 6g cross. The difference across actions is greater than within each action (table S1, ref. 1.12,1.13). (**F-H**) Data are mean+s.e.m. over n=41 sessions from 4 mice.

Mice performed isometric actions in order to receive sucrose reinforcement. We defined two targets as push and pull that exceed thresholds of force and duration determined for each session (Fig. 1B,C, Table 2,3, Methods), and these actions triggered delayed sucrose reward (5 µl per reward). Mice selectively increased performance of whichever action was reinforced, confirming that they performed actions to obtain sucrose (Fig. 1D, S2A-C, Methods).

We characterized the simultaneous activity of four forelimb muscles - biceps, triceps, palmaris longus (PL), and the extensor digitorium communis (EDC) - during the performance of these isometric actions (Fig. 1E). Isometric push and pull were executed with co-contraction of flexors (biceps, PL) and extensors (triceps, EDC) (Fig. 1F,G), but the pattern across muscles was action-specific (Fig. 1H). Thus, isometric push and pull are highly granular actions, distinguished not by muscle recruitment but by subtle differences in co-contraction patterns within the same forelimb.

### Striatal ensembles encode the identity of isometric actions

To test whether the striatum encodes the identity of isometric forelimb actions, we measured the simultaneous activity of D1- and D2-SPNs in the forelimb region of dorsolateral striatum (*48, 49*) (DLS) (Fig. 2A,B). To identify D1- and D2-SPNs within the same subject (*50*), we used transgenic mice that expressed the red label tdTomato either in D1-SPNs (T6(*51*) and Ai9(*52*) x EY217(*53*)) or D2-SPNs (Ai9(*52*) x Adora2cre(*53*)) (Table 1), and we developed an automated method to classify red versus non-red neurons (fig. S1). tdTomato-positive neurons were thus identified as D1-SPNs (or D2-SPNs, depending on the mouse line), and tdTomato-negative neurons were classified as the other subtype; together, D1- and D2-SPNs account for approximately 95% of all striatal neurons, so non-red neurons are overwhelmingly the other SPN subtype (*50, 54, 55*). We virally expressed GCaMP6f in both SPN populations in DLS using the *Camk2a* promoter (*50*) and used 2-photon microscopy through a 1mm diameter GRIN lens to image neuronal calcium activity as a proxy for action potentials. We were able to image up to ∼200 neurons simultaneously, and the fluorescence signal in both color channels was stable across months of animal training (fig. S3K). We identified ongoing isometric actions that exceeded 3g of force for ∼66 ms duration (two imaging frames) and that were temporally isolated (i.e. had no preceding action in the 0.5 seconds before onset). The activity of D1- and D2-SPNs was modulated for both actions (Fig. 2B,C), with slightly greater activity in D1-SPNs (Fig. 2C).

**Fig. 2.**
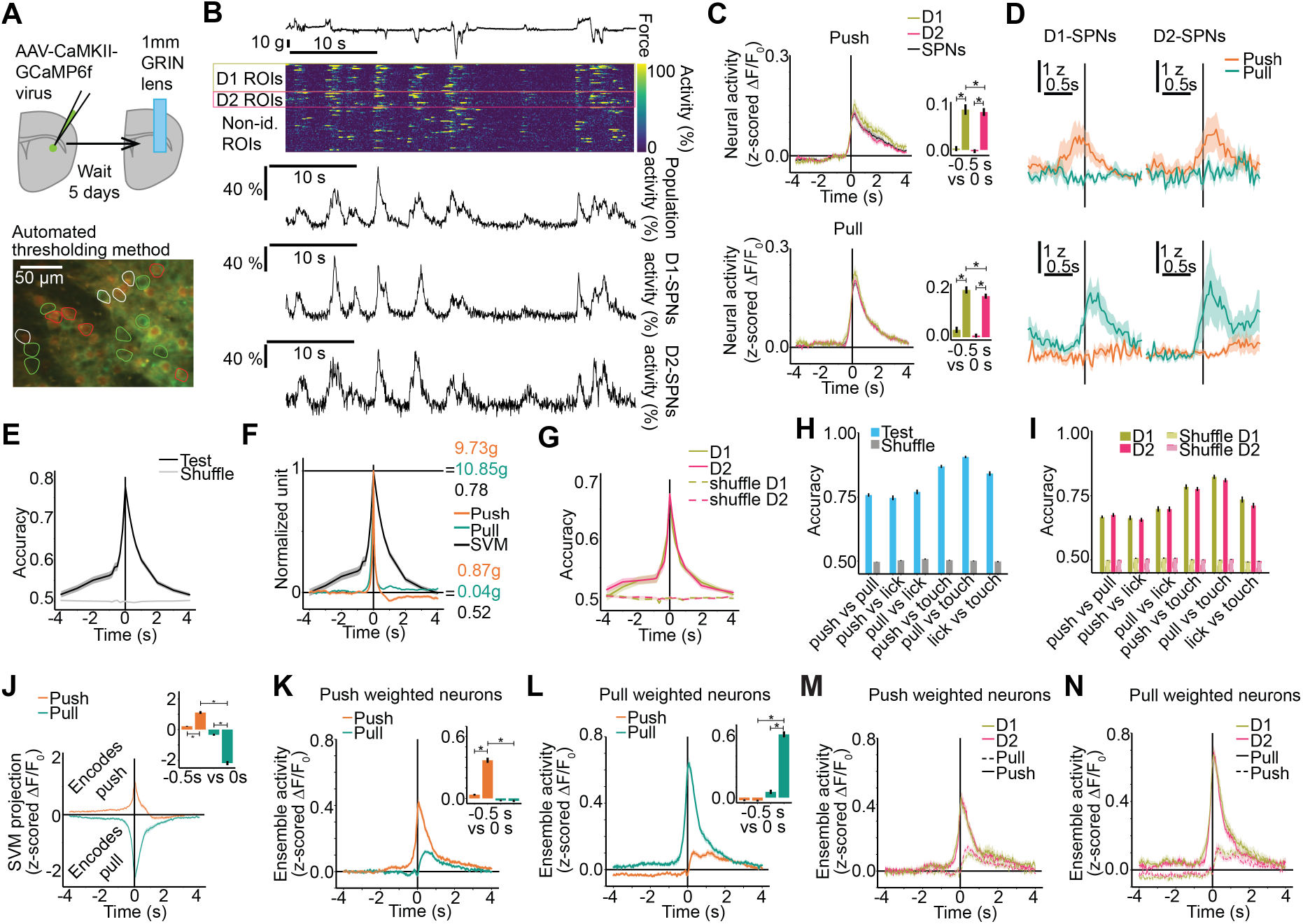
D1- and D2-SPNs both encode identity of isometric forelimb actions. (**A**) Schematic of experimental procedures. Top: GCaMP6f virus injection and lens implant in transgenic mice to image D1- and D2-SPNs simultaneously. Bottom: Schematic of automated method for identifying D1- and D2-SPNs based on transgenic expression of tdTomato. (**B**) Example trace of force, individual neuronal activity for all ROIs in this field of view, population activity averaged across neurons, averaged activity for D1-SPNs (n=21 neurons) and averaged activity for D2-SPNs (n=21 neurons). Neural activity is ΔF/F_0_. (**C**) Population activity locked to peak force of push and pull actions, averaged across D1-, D2-, and all SPNs. Data are mean+s.e.m. across 86 sessions from 8 mice. Activity increased at push force peak (“0 s”) relative to “-0.5 s” for D1-SPNs (table S1, ref. 2.1) and D2-SPNs (table S1, ref. 2.2). Activity increased at pull force peak (“0 s”) relative to “-0.5 s” for D1-SPNs (table S1, ref. 2.3) and D2-SPNs (table S1, ref. 2.4). D1-SPNs were more active than D2-SPNs at force peak (table S1, ref. 2.5,2.6). Bar plots were ΔF/F_0_ averaged across 3 event-centered imaging frames. (**D**) Example traces of individual D1- or D2-SPN activity locked to force peak for push and pull actions. (**E**) Decoding all SPN activity to predict action identity on single trials with a model fit at each time lag relative to force peak. Accuracy at force peak was greater than a model (“shuffle”) trained with shuffled action identity (table S1, ref. 2.7). Accuracy was greater than shuffle when averaged over time points in the 4 seconds before force peak (table S1, ref. 2.8) and in the 0.5 seconds before force peak during which there was no other action (table S1, ref. 2.9). Data are mean+s.e.m. across n=87 sessions from 8 mice. (**F**) Decoding all SPN activity overlayed on the average force of actions (normalized to 0 at -4s and 1 at peak of 0s). (**G**) Decoding D1- or D2-SPN activity separately. Accuracy at force peak was greater than shuffle for D1-SPNs (table S1, ref. 2.10) and D2-SPNs (table S1, ref. 2.11), with no significant difference between D1- and D2-SPNs (table S1, ref. 2.12). Accuracy was greater than shuffle when averaged over time points in the 4 seconds before force peak for D1-SPNs (table S1, ref. 2.13) and D2-SPNs (table S1, ref. 2.14) and in the 0.5 seconds before force peak for D1-SPNs (table S1, ref. 2.15) and D2-SPNs (table S1, ref. 2.16). Data are mean+s.e.m. across n=87 sessions from 8 mice. (**H-L**) Data are mean+s.e.m. across 86 sessions from 8 mice. (**H**) Decoding SPN activity at force peak for push and pull, predicting action identity between pairs of actions including push, pull, touch, and licking. Accuracy was greater than shuffle for all action pairs (table S1, ref. 2.17-2.22). (**I**) Same as (G), decoding D1- and D2-SPNs activity. Decoding push versus pull was not significantly different between D1- and D2-SPNs (table S1, ref. 2.23). Accuracy was greater than shuffle for all action pairs for D1-SPNs (table S1, ref. 2.24-2.29) and D2-SPNs (table S1, ref. 2.30-2.35). (**J**) Projection of population activity (z-scored ΔF/F_0_) on the SVM dimension locked to force peak. Inset: average SVM projection at 500 ms before force peak (“-0.5 s”) and at force peak (“0 s”). For the push action, the SVM projection at force peak (“0 s”) increased relative to “-0.5 s” (table S1, ref. 2.36). For the pull action, the SVM projection at force peak (“0 s”) decreased relative to “-0.5 s” (table S1, ref. 2.37). The SVM projection at force peak was greater for push than for pull (table S1, ref. 2.38). (**K**) Ensemble activity (z-scored ΔF/F_0_) averaged across push weighted neurons, locked to force peak for push and pull actions. Inset: average activity at -0.5 s before force peak and at force peak for each action. The activity of push weighted neurons increased at push force peak (“0 s”) relative to “-0.5 s” (table S1, ref. 2.39). Activity at push force peak was greater than at pull force peak (table S1, ref. 2.40). (**L**) Ensemble activity (z-scored ΔF/F_0_) averaged across pull weighted neurons, locked to force peak for push and pull actions. Inset: average activity at -0.5 s before force peak and at force peak for each action. The activity of pull weighted neurons increased at pull force peak (“0 s”) relative to “-0.5 s” (table S1, ref. 2.41). Activity at pull force peak was greater than at push force peak (table S1, ref. 2.42). (**M**) Same as (K) for D1- and D2-SPNs. Activity of push weighted D1- and D2-SPNs around push force peak (-0.5 s to 0.5 s) is not significantly different (table S1, ref. 2.43). Data are mean+s.e.m. across 63 sessions from 8 mice. (**N**) Same as (L) for D1- and D2-SPNs. Activity of pull weighted D1- and D2-SPNs around pull force peak (-0.5 s to 0.5 s) is not significantly different (table S1, ref. 2.44). Data are mean+s.e.m. across 73 sessions from 8 mice.

Many individual neurons modulated their activity only during a specific action (Fig. 2D). We therefore asked whether the spatiotemporal activity of SPNs encoded the identity of ongoing actions. We extracted the vector of SPN population activity averaged in 150 ms bins (5 imaging frames) centered at time lags relative to force peak for each trial, and we decoded single-trial population activity into action identity (push or pull) using a linear support vector machine (training in 90% of data, testing in 10% held out data). For time lags before (after) force peak, we analyzed trials with no other action in the 0.5 seconds before (after) force peak. Population activity predicted action identity well above chance with peak accuracy at the time of peak force (Fig. 2E,F). In addition, population activity predicted action identity in advance of force peak (Fig. 2F). Interestingly, both D1- and D2- SPNs encoded action identity with equal accuracy and timing (Fig. 2G).

Population activity at force peak predicted action identity even for trials when other actions were allowed in the 0.5s temporal window before and after (Fig. 2H). D1- and D2-SPNs encoded these actions with equal accuracy (Fig. 2I). Given that the striatum encodes continuous kinematics (*22, 23*), we verified action identity was encoded (fig. S3I,J) for trials with matched continuous parameters (fig. S3A-H) including force, duration, and licking.

To understand the structure of this encoding, we examined how action identity was embedded within the high dimensional space of population activity where each neuron’s instantaneous activity is one axis. The SVM identifies a single dimension that best predicts the animal’s action at the time of the force peak. It follows that the projection of population activity on that SVM dimension (a weighted sum over neurons) showed differential modulation at force peak for push and pull actions (Fig. 2J). The SVM’s weights partition neurons into action-specific groups whose activation biases the decoder to predict push or pull. The push-weighted neurons increased activity at push force peak, but not for pull force peak (Fig. 2K). Similarly, the pull weighted neurons increased activity at pull force peak but not for push force peak (Fig. 2L). D1- and D2-SPNs showed no difference in SVM weights and action-specific modulation of push- and pull-weighted neurons (Fig. 2M,N).

The striatum has a classical role in the reinforcement of actions, with D1- and D2-SPNs thought to have opponent (*6, 12*) or different (*13*) roles. Therefore, we examined if encoding of action identity would change whether the action is reinforced or not (fig. S2A-C). We found that both types of SPNs encode ongoing action identity regardless of whether both or one of the particular actions were reinforced (fig. S2D-G). Further, we tested whether striatal plasticity is necessary for action encoding. Notably, we found that striatal activity encoded the identity of specific actions even in mice with impaired striatal plasticity (*56*) who could not learn which actions were reinforced (*57–59*) (fig. S2H-J). These results reveal that although the striatum has a clear role in reinforcement and learning, it can encode ongoing actions independently of these roles.

Together, these results demonstrate that both D1- and D2-SPNs robustly encode the identity of ongoing actions at a fine-grained level.

### Closed-loop stimulation of action-specific ensembles

We next addressed the critical question of whether these ensembles causally control the specific actions they encode. One technical challenge was that we needed to target perturbations to specific and spatially-intermingled D1- and D2- SPNs that encode action identity. For this, we developed an approach to stimulate individual SPNs with 2-photon optogenetics through a GRIN lens (*41*) in mice co-expressing GCaMP6f and red-shifted activating opsin ChRmine (*40*) (Fig. 3A-C). We used a spatial light modulator to holographically target a group of user-defined SPNs for simultaneous stimulation (*39*). Another technical challenge was that action duration was very short, and we needed to trigger stimulation upon self-initiated actions with low latency, as force onsets rapidly over milliseconds (Fig. 1C). For this, we developed an experimental platform that allowed us to flexibly select the trigger action and stimulation pattern and begin the stimulation with ∼2ms total latency (Fig. 3D and S4B).

**Fig. 3.**
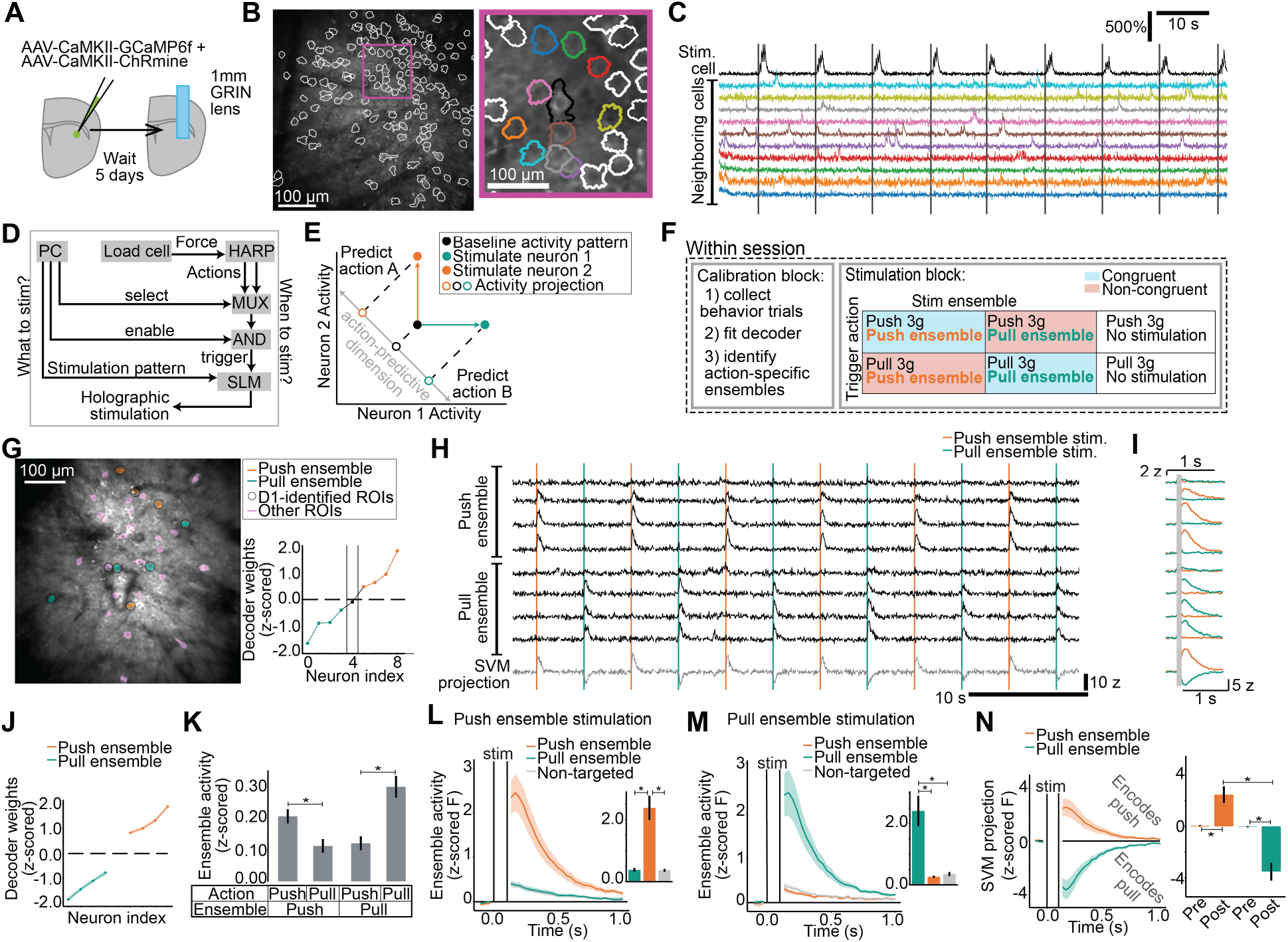
Closed-loop modeling and manipulation of neuronal ensembles that encode action identity. **(A**) Schematic of GCaMP6f and ChRmine viral injection and lens implant in transgenic mice to image and photostimulate D1- and D2-SPNs. (**B**) Example field of view with a single neuron targeted for stimulation. Inset: zoom in onto targeted neuron in black. (**C**) Example activity traces of neurons in (B) inset. Targeted neuron (in black) and neighboring non-targeted neurons’ activity during 2-photon optogenetic stimulation. (**D**) Schematic of closed-loop system to select the pattern of holographic stimulation and the action that triggers stimulation. (**E**) Cartoon depicting how activating specific neurons can manipulate a dimension of population activity that encodes and predicts specific actions. (**F**) Schematic for conducting the modeling and manipulation experiment. (**G**) Left: example field of view with regions of interest labeled based on their decoder weights and identification as D1-SPNs. Right: Decoder weights for identified D1-SPNs, colored by assignment into action-specific ensembles. (**H**) Z-scored fluorescence of each ROI member of each action-specific ensemble, and the projection of all population activity on the SVM decoder dimension. Four consecutive stimulation trials concatenated from the beginning, middle, and end of the example session. (**I**) Trial-averaged traces of targeted neurons and projection of population activity on the action-predictive SVM dimension. (**J-N**) Data are mean+s.e.m. across n=21 sessions and 10 mice. (**J**) Average decoder weights of action-specific ensembles. The top four neurons with the largest weights are shown. (**K**) Average activity (z-scored fluorescence) of action-specific ensembles in 100 ms before force peak during trials in the calibration block. Push ensemble activity was larger for push than pull (table S1, ref. 3.1). Pull ensemble activity was larger for pull than push (table S1, ref. 3.2). (**L-M**) Neural activity averaged across neurons (z-scored fluorescence) within each ensemble, locked to stimulation of each ensemble. Inset: average ensemble activity in 100 ms after stimulation. (**L**) After push ensemble stimulation, activity of the push ensemble was greater than the pull ensemble (table S1, ref. 3.3) and non-targeted neurons (table S1, ref. 3.4). There was no significant difference between the pull ensemble and non-targeted neurons (table S1, ref. 3.5). (**M**) After pull ensemble stimulation, activity of the pull ensemble was greater than the push ensemble (table S1, ref. 3.6) and non-targeted neurons (table S1, ref. 3.7). There was no significant difference between the push ensemble and non-targeted neurons (table S1, ref. 3.8). (**N**) Projection of population activity on the SVM dimension locked to ensemble stimulation. Inset: Average SVM projection in the 100 ms before stimulation (“pre”) and after stimulation (“post”). Stimulating the push ensemble increased the “post” SVM projection relative to “pre” (table S1, ref. 3.9). Stimulating the pull ensemble decreased the “post” SVM projection relative to “pre” (table S1, ref. 3.10). The “post” SVM projection was greater after push ensemble stimulation than pull ensemble stimulation (table S1, ref. 3.11).

Within a single session, we identified action-specific SPN ensembles and stimulated them in closed-loop during specific, self-paced actions (Fig. 3E,F). In a calibration block, we collected behavior trials and fit a decoder to predict action identity from SPN activity. As before (Fig. 2K-N), we identified action-specific ensembles (Fig. 3G) using the weights of the force peak SVM. We defined the action-specific ensembles as equally sized groups of 4-11 neurons with the largest decoder weights (mean ensemble size was 6.7 neurons over 21 sessions). Then in the stimulation block, specific self-paced actions triggered stimulation of action-specific ensembles (Fig. 3H,I). We interleaved 6 conditions: two actions that triggered stimulation (3g push, 3g pull) and three stimulation patterns for each action (push ensemble, pull ensemble, no stimulation) (Fig. 3F).

The identified ensembles had similar magnitude decoder weights (Fig. 3J) and modulated specifically for their corresponding actions (Fig. 3K). Stimulation specifically activated the targeted ensemble (Fig. 3L,M). We verified that this stimulation of a small number of action-specific neurons bi-directionally drove the action-specific SVM dimension of population activity (Fig. 3N). Thus, selective stimulation of small, action-specific striatal ensembles was sufficient to bias population activity toward encoding the corresponding action (Fig. 3E).

### Specific striatal ensembles control specific actions

We used this approach to test whether action-specific striatal ensembles control specific actions or simply invigorate any ongoing action. If ensembles control specific actions, stimulating a pull ensemble should selectively increase pull force, with no effect or an opposing effect during push. If ensembles merely invigorate movement, any ensemble stimulation should increase force regardless of action identity. We refer to the pairing of an ongoing action with ensemble stimulation specific to that action as “congruent” (e.g. pull ensemble stimulation during pull action), and the pairing of an ongoing action with ensemble stimulation specific to the other action as “non-congruent” (e.g. push ensemble stimulation during pull action) (Fig. 3F). This also allowed us to test whether D1-SPNs promote and D2-SPNs suppress action, or both promote action.

Stimulating specific D1-SPN ensembles congruent with the ongoing action increased ongoing force (Fig. 4A left). In contrast, no change in ongoing force was observed after stimulating the non-congruent ensemble (Fig. 4A right). Notably, stimulating specific D2-SPN ensembles congruent with the ongoing action also increased ongoing force (Fig. 4B left), whereas stimulation of the non-congruent ensemble did not (Fig. 4B right). In other words, stimulation of the same ensemble during different ongoing actions had different effects (e.g. stimulating the pull ensemble increased force during pull, but not during push).

**Fig. 4.**
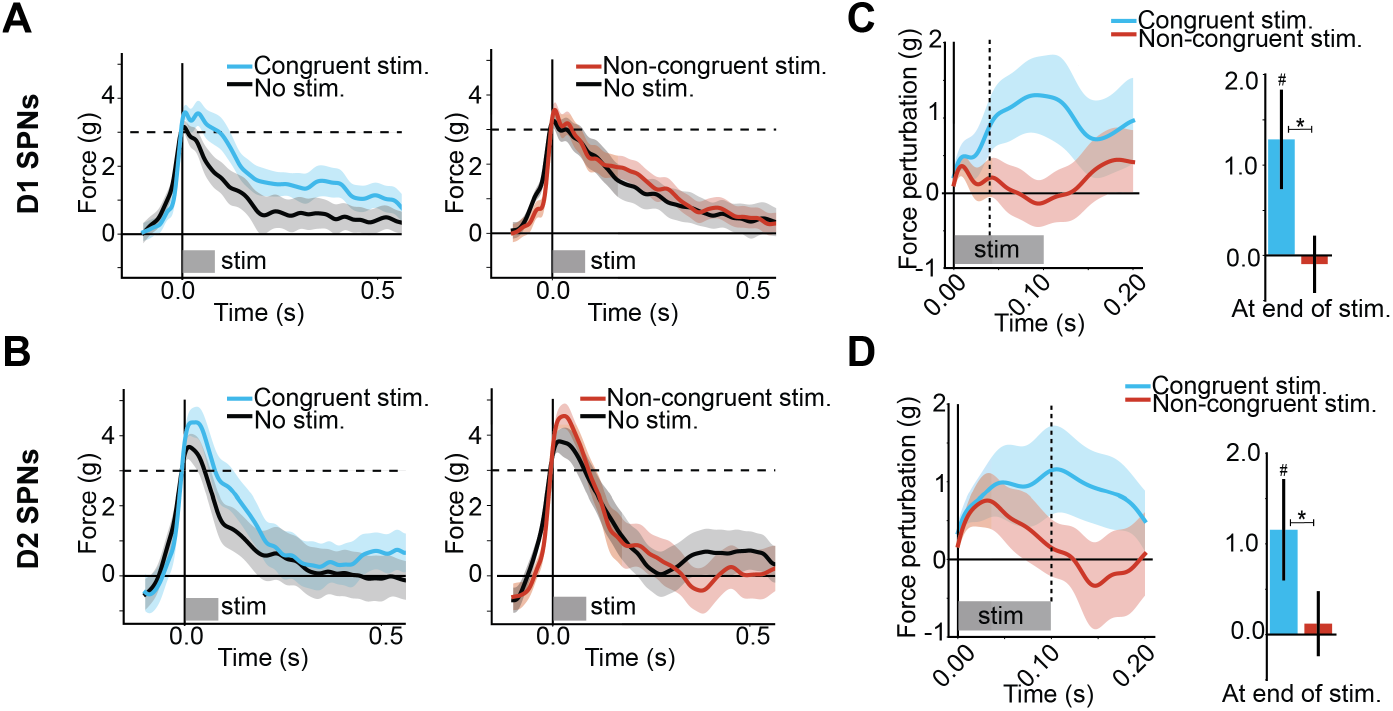
Action-specific ensembles of D1- and D2-SPNs control congruent actions. (**A-B**) Trial-averaged force traces locked to 3g cross that triggers holographic stimulation. In this plotting convention, force is positive for the action executed at time 0 (push or pull) and negative for the other action. (**A**) Stimulation of D1-SPN ensembles. Data are mean+s.e.m. across n=12 sessions and 9 mice. (**B**) Stimulation of D2-SPN ensembles. Data are mean+s.e.m. across n=9 sessions and 5 mice. (**C-D**) Force perturbation, i.e. the trial-averaged force trace on stimulation trials minus non-stimulation trials. Right panel is the force perturbation at the end of stimulation, averaged from 100ms to 110ms. (**C**) Force perturbation from stimulation of D1-SPN ensembles. The perturbation was greater than 0 with congruent stimulation (table S1, ref. 4.12) and was greater than non-congruent stimulation (table S1, ref. 4.11) at the end of stimulation. The perturbation was not significantly different than 0 at the end of non-congruent stimulation (table S1, ref. 4.13). The perturbation first became significantly different between congruent and non-congruent stimulation at 40ms after stimulation onset (table S1, ref. 4.5). For congruent stimulation, the perturbation was significantly different from 0 in the intervals from 0-50ms (table S1, ref. 4.27), 50-100ms (table S1, ref. 4.28), and 100-150ms (table S1, ref. 4.29). For non-congruent stimulation, the perturbation was not significantly different from 0 in those same intervals (table S1, ref. 4.30, 4.31, and 4.32). (**D**) Force perturbation from stimulation of D2-SPN ensembles. The perturbation was greater than 0 with congruent stimulation (table S1, ref. 4.25) and was greater than non-congruent stimulation (table S1, ref. 4.24) at the end of stimulation. The perturbation was not significantly different than 0 at the end of non-congruent stimulation (table S1, ref. 4.26). The perturbation first became significantly different between congruent and non-congruent stimulation at 100ms after stimulation onset (table S1, ref. 4.24). For congruent stimulation, the perturbation was significantly different from 0 in the intervals from 0-50ms (table S1, ref. 4.33), 50-100ms (table S1, 4.34), and 100-150ms (table S1, 4.35). For non-congruent stimulation, the perturbation was significantly different from 0 in the interval 0-50ms (table S1, ref. 4.36), but not in the intervals from 50-100ms (table S1, ref. 4.37) and 100-150ms (table S1, ref. 4.38).

We quantified the force perturbation as the difference between force traces in stimulation trials and non-stimulation trials. Stimulation of action-specific ensembles increased congruent but not non-congruent force for both D1- and D2-SPNs (Fig. 4C,D). Congruent stimulation of D1-SPN ensembles increased perturbation relative to non-congruent stimulation as early as 40 ms after start of stimulation, whereas D2-SPNs ensemble stimulation increased perturbation later, around 100 ms after start of stimulation. Indeed, D2-SPNs transiently increased force in the first 50ms of stimulation for both congruent and non-congruent conditions, followed by a selective increase in force for the congruent condition. Thus, the perturbation dynamics of D1- vs D2-SPNs were slightly different, suggesting that D1 and D2 populations enable different execution dynamics potentially through different downstream circuits. Together, these results demonstrate that specific ensembles of both D1- and D2-SPNs causally control specific ongoing actions, selectively increasing force only when stimulation was congruent with the ongoing action. Rather than simply invigorating movement, striatal ensembles bias specific actions in real time.

## Discussion

Our results reveal that the striatum operates at a previously unappreciated level of granularity, specifying actions as fine as distinct patterns of muscle co-contraction within the same limb. While prior work has shown that striatal activity encodes movements, our findings demonstrate that small, spatially intermingled ensembles of striatal neurons causally control ongoing actions in real time. This establishes the striatum not as a structure that merely invigorates or reinforces movement, but as a system that specifies and biases action execution. Importantly, this control is highly specific: stimulation of action-specific ensembles selectively increases force only when matched to the ongoing action, ruling out a generalized increase in movement vigor. Furthermore, our findings show that both D1- and D2-SPNs contribute similarly to the specification and control of actions. This challenges the classical framework supported mainly through bulk manipulations, assigning opponent roles to these populations in movement promotion and suppression. Instead, our findings support a framework in which specific populations of D1- and D2-SPNs participate in moment-to-moment (*22, 23*) fine motor control.

These experiments reveal that the population dynamics of D1- and D2-SPNs encode action with high resolution in movement as well as time. The task employed here enabled us to test whether the striatum can encode different self-paced forelimb actions with similar force and no overt kinematics, just different patterns of forelimb muscle activation. Further, at each moment of execution, different population activity patterns specified action. Decoders trained at time lags locked to 3g cross significantly predicted action identity (table S1, ref 2.45). The dimension of population activity that encoded action identity was different across time lags (table S1, ref 2.46-2.53), and became more aligned with the force peak SVM with time (table S1, ref 2.46). Thus, a given ensemble is not a labelled line for action with simple ramping activity; instead population activity dynamically specifies action from moment-to-moment. These results for isometric actions are consistent with recent work showing striatal activity encodes moment-to-moment kinematics of overt movement (*22, 23*).

These results also offer new views on the canonical model that D1- and D2-SPNs promote and suppress movement respectively (*16, 25, 28, 29*). The fact that both D1 and D2-SPNs equally encode action identity is consistent with previous work showing that, despite having distinct cortical inputs, both cell types receive similar motor information (*32*).

Furthermore, whereas bulk manipulations have supported opposing effects (*6, 28*), subsequent manipulations (*36, 37*) suggested that their coordinated activity are necessary for movement execution. Here we identified and stimulated small action-specific ensembles of D1- and D2-SPNs and found that both increase the force of congruent actions. Thus, D2-SPNs can not only promote movement, but also control specific actions in real time. This finding suggests that D2-SPN projections to GPe can have effects that are independent of the canonical GPe-mediated increase in activity of GPi/SNr (*60*). Rather, our results are consistent with other work from our team showing that GPe projects directly to cortex, thalamus and brainstem, and that D2-SPNs can therefore disinhibit movement through the inhibition of specific GPe projections that inhibit thalamus and brainstem (*61*). Further, that study shows that GPe modulates forelimb movement through the parafascicular thalamus. Thus, D1- and D2-SPNs could have different output circuits that affect the same forelimb action. The fact that D1- and D2-SPN stimulation produced different perturbation dynamics suggests that they not only act through distinct descending circuits, but may also serve different roles in execution. It is also important to note that we focused our recordings and manipulations on forelimb region of dorsolateral striatum, and understanding the function of D1- and D2-SPNs would likely depend on the striatal subregion (*22, 29, 62*), each receiving distinct topographic inputs from cortex (*48, 49*) and thalamus, and also projecting to specific outputs.

This study positions the striatum as a center for controlling granular ongoing actions, despite its indirect connections to the spinal cord. Although basal ganglia outputs tonically suppress downstream motor centers (*24, 63–65*), recent work reveals highly specific output dynamics during forelimb behavior (*66*). Our work provides a foundation for future work on a fundamental problem: to understand how action-specific ensembles in the center of the brain can have such remarkably specific control over downstream motor centers. Conceptually, we hypothesize that the striatum guides motor centers into specific subspaces of neural population dynamics which implement action plans and execution (*67*), consistent with evidence that striatal activity (*68*) and corticostriatal plasticity (*57*) are required for the brain to learn and control low-dimensional cortical dynamics (*69–71*) that directly operate brain-machine interfaces. More generally, our experimental approach can be applied to motor circuits across the brain to test not only what neurons encode but also what their activity causally produces (*72*), both in terms of downstream population dynamics and movement.

The discovery that striatal ensembles control granular actions offers a framework for understanding movement disorders with striatal dysfunction characterized by impaired, highly specific motor pattern, such as task-specific dystonias (*27*) (e.g. writer’s cramp, musician’s dystonia) and the choreic disruptions of selective movements observed in Huntington’s disease (*15, 25*).

## Acknowledgments

We thank C. Maos de Ferro for development of load sensor hardware and software, and the members of the Scientific Hardware Platform of the Champalimaud Institute for help with hardware and software. We thank T. Tabachnik and R. Hormigo for technical advice and hardware support. We thank A. Vaz, M. Correia and G. Martins for mouse colony management. We thank L. Hammond and the Zukerman Institute’s Cellular Imaging Platform for guidance in imaging and analysis. We thank A.M. Vicente for blinding the experimenter and discussions about mouse training. We thank Bruker and M.J. Fox for help developing the Bruker Software API. We thank K. Deisseroth, Y.S. Kim, and C. Ramakrishnan for providing the ChRmine virus for holographic stimulation. During the preparation of this work, the authors used Claude to improve text readability and compress word count. After using this tool, the authors reviewed and edited the content as needed and take full responsibility for the content of the publication.

## Funding

Fundacao para Ciencia e Tecnologia grant FCT PD/BD/105950/2014 (IR-V) BRAIN Initiative National Institute of Mental Health postdoctoral fellowship 1F32MH118714-01 (VRA)

NIH NINDS Pathway to Independence Award 1K99NS128250-01(VRA)

Aligning Science Across Parkinson’s (ASAP-020551) through the Michael J. Fox Foundation for Parkinson’s Research (MJFF) (IR-V, VRA, DSP, RMC)

NS NS104649 (RMC)

Simons-Emory International Consortium Simons 717104 (RMC)

National Institutes of Health 5U19NS104649 (DSP and RMC)

## Author contributions

IR-V, VRA, DSP and RMC designed the study, interpreted results and wrote the manuscript. IR-V, VRA, HFMR, DSP, and RMC developed experimental setups. IR-V performed experiments. IR-V and VRA analyzed data and developed the model-guided system for closed-loop stimulation.

## Competing interests

DSP is listed as an inventor of the following patent: “Devices, apparatus and method for providing photostimulation and imaging of structures” (United States Patent 9846313). He also receives income as a consultant to the Bruker Corporation related to the above patent. IR-V, VRA, HFMR, and RMC declare no competing interests.

## Data, code, and materials availability

All data and materials used for this work are described in table S6, where *doi*s for public repository are available. Data is available in the BioStudies repository: 10.6019/S-BIAD3134 and 10.6019/S-BIAD3135. Code is available on Zenodo: 10.5281/zenodo.17821678.

## Supplementary Materials

Materials and Methods

Supplementary Text

Figures S1 to S4

Tables S1 to S6

References (73-79)

## Notes

### Summary of Updates

In the last version update we updated the title of the manuscript but not in the submission so the tittle was different between the manuscript and the bioRxiv website.

## References and Notes

1. S. Arber, R. M. Costa, Connecting neuronal circuits for movement. Science 360, 1403–1404 (2018).

2. P. Mazzoni, A. Hristova, J. W. Krakauer, Why Don’t We Move Faster? Parkinson’s Disease, Movement Vigor, and Implicit Motivation. J. Neurosci. 27, 7105 (2007).

3. B. Panigrahi, K. A. Martin, Y. Li, A. R. Graves, A. Vollmer, L. Olson, B. D. Mensh, A. Y. Karpova, J. T. Dudman, Dopamine Is Required for the Neural Representation and Control of Movement Vigor. Cell 162, 1418–1430 (2015).

4. J. T. Dudman, J. W. Krakauer, The basal ganglia: from motor commands to the control of vigor. Current Opinion in Neurobiology 37, 158–166 (2016).

5. R. S. Turner, M. Desmurget, Basal ganglia contributions to motor control: a vigorous tutor. Current Opinion in Neurobiology 20, 704–716 (2010).

6. E. A. Yttri, J. T. Dudman, Opponent and bidirectional control of movement velocity in the basal ganglia. Nature 533, 402–406 (2016).

7. D. Joel, Y. Niv, E. Ruppin, Actor–critic models of the basal ganglia: new anatomical and computational perspectives. Neural Networks 15, 535–547 (2002).

8. Y. Niv, Reinforcement learning in the brain. Journal of Mathematical Psychology 53, 139–154 (2009).

9. M. Ito, K. Doya, Multiple representations and algorithms for reinforcement learning in the cortico-basal ganglia circuit. Current Opinion in Neurobiology 21, 368–373 (2011).

10. K. Samejima, Y. Ueda, K. Doya, M. Kimura, Representation of Action-Specific Reward Values in the Striatum. Science 310, 1337–1340 (2005).

11. J. N. J. Reynolds, B. I. Hyland, J. R. Wickens, A cellular mechanism of reward-related learning. Nature 413, 67–70 (2001).

12. A. V. Kravitz, L. D. Tye, A. C. Kreitzer, Distinct roles for direct and indirect pathway striatal neurons in reinforcement. Nature Neuroscience 15, 816–818 (2012).

13. A. M. Vicente, P. Galvão-Ferreira, F. Tecuapetla, R. M. Costa, Direct and indirect dorsolateral striatum pathways reinforce different action strategies. Current Biology 26, R267–R269 (2016).

14. I. Donaldson, C. D. Marsden, S. Schneider, K. Bhatia, Marsden’s Book of Movement Disorders (Oxford University Press, 2012; 10.1093/med/9780192619112.001.0001).

15. R. L. Albin, A. B. Young, J. B. Penney, The functional anatomy of basal ganglia disorders. Trends in Neurosciences 12, 366–375 (1989).

16. M. R. DeLong, Primate models of movement disorders of basal ganglia origin. Trends in Neurosciences 13, 281–285 (1990).

17. N. Giladi, D. McMahon, S. Przedborski, E. Flaster, S. Guillory, V. Kostic, S. Fahn, Motor blocks in Parkinson’s disease. Neurology 42, 333–333 (1992).

18. D. Shohamy, C. E. Myers, S. Grossman, J. Sage, M. A. Gluck, R. A. Poldrack, Cortico-striatal contributions to feedback-based learning: converging data from neuroimaging and neuropsychology. Brain 127, 851–859 (2004).

19. B. J. Knowlton, J. A. Mangels, L. R. Squire, A Neostriatal Habit Learning System in Humans. Science 273, 1399–1402 (1996).

20. A. Klaus, G. J. Martins, V. B. Paixao, P. Zhou, L. Paninski, R. M. Costa, The Spatiotemporal Organization of the Striatum Encodes Action Space. Neuron 95, 1171–1180.e7 (2017).

21. J. E. Markowitz, W. F. Gillis, C. C. Beron, S. Q. Neufeld, K. Robertson, N. D. Bhagat, R. E. Peterson, E. Peterson, M. Hyun, S. W. Linderman, B. L. Sabatini, S. R. Datta, The Striatum Organizes 3D Behavior via Moment-to-Moment Action Selection. Cell 174, 44–58.e17 (2018).

22. A. K. Dhawale, S. B. E. Wolff, R. Ko, B. P. Ölveczky, The basal ganglia control the detailed kinematics of learned motor skills. Nature Neuroscience 24, 1256–1269 (2021).

23. J. Park, P. Polidoro, C. Fortunato, J. Arnold, B. Mensh, J. A. Gallego, J. T. Dudman, Conjoint specification of action by neocortex and striatum. Neuron 113, 620–636.e6 (2025).

24. O. Hikosaka, Y. Takikawa, R. Kawagoe, Role of the Basal Ganglia in the Control of Purposive Saccadic Eye Movements. Physiological Reviews 80, 953–978 (2000).

25. R. L. Albin, A. Reiner, K. D. Anderson, J. B. Penney, A. B. Young, Striatal and nigral neuron subpopulations in rigid Huntington’s disease: Implications for the functional anatomy of chorea and rigidity-akinesia. Annals of Neurology 27, 357–365 (1990).

26. K. Simonyan, H. Cho, A. Hamzehei Sichani, E. Rubien-Thomas, M. Hallett, The direct basal ganglia pathway is hyperfunctional in focal dystonia. Brain 140, 3179–3190 (2017).

27. C. M. Stahl, S. J. Frucht, Focal task specific dystonia: a review and update. J Neurol 264, 1536–1541 (2017).

28. A. V. Kravitz, B. S. Freeze, P. R. L. Parker, K. Kay, M. T. Thwin, K. Deisseroth, A. C. Kreitzer, Regulation of parkinsonian motor behaviours by optogenetic control of basal ganglia circuitry. Nature 466, 622–626 (2010).

29. B. F. Cruz, G. Guiomar, S. Soares, A. Motiwala, C. K. Machens, J. J. Paton, Action suppression reveals opponent parallel control via striatal circuits. Nature 607, 521–526 (2022).

30. M. T. Kaufman, M. M. Churchland, S. I. Ryu, K. V. Shenoy, Cortical activity in the null space: permitting preparation without movement. Nature Neuroscience 17, 440–448 (2014).

31. H. K. Inagaki, S. Chen, K. Daie, A. Finkelstein, L. Fontolan, S. Romani, K. Svoboda, Neural Algorithms and Circuits for Motor Planning. Annual Review of Neuroscience 45, 249–271 (2022).

32. A. Nelson, B. Abdelmesih, R. M. Costa, Corticospinal populations broadcast complex motor signals to coordinated spinal and striatal circuits. Nature Neuroscience 24, 1721–1732 (2021).

33. M. N. Economo, S. Viswanathan, B. Tasic, E. Bas, J. Winnubst, V. Menon, L. T. Graybuck, T. N. Nguyen, K. Smith, Z. Yao, L. Wang, C. R. Gerfen, J. Chandrashekar, H. Zeng, L. L. Looger, K. Svoboda, Distinct descending motor cortex pathways and their roles in movement. Nature 563, 79–84 (2018).

34. M. M. Churchland, K. V. Shenoy, Preparatory activity and the expansive null-space. Nature Reviews Neuroscience 25, 213–236 (2024).

35. G. Cui, S. B. Jun, X. Jin, M. D. Pham, S. S. Vogel, D. M. Lovinger, R. M. Costa, Concurrent activation of striatal direct and indirect pathways during action initiation. Nature 494, 238–242 (2013).

36. M. Sheng, D. Lu, Z. Shen, M. Poo, Emergence of stable striatal D1R and D2R neuronal ensembles with distinct firing sequence during motor learning. Proceedings of the National Academy of Sciences 116, 11038–11047 (2019).

37. F. Tecuapetla, X. Jin, S. Q. Lima, R. M. Costa, Complementary Contributions of Striatal Projection Pathways to Action Initiation and Execution. Cell 166, 703–715 (2016).

38. Z. Zhang, L. E. Russell, A. M. Packer, O. M. Gauld, M. Häusser, Closed-loop all-optical interrogation of neural circuits in vivo. Nature Methods 15, 1037–1040 (2018).

39. W. Yang, L. Carrillo-Reid, Y. Bando, D. S. Peterka, R. Yuste, Simultaneous two-photon imaging and two-photon optogenetics of cortical circuits in three dimensions. eLife 7, 1–21 (2018).

40. J. H. Marshel, Y. S. Kim, T. A. Machado, S. Quirin, B. Benson, J. Kadmon, C. Raja, A. Chibukhchyan, C. Ramakrishnan, M. Inoue, J. C. Shane, D. J. McKnight, S. Yoshizawa, H. E. Kato, S. Ganguli, K. Deisseroth, Cortical layer–specific critical dynamics triggering perception. Science 365, eaaw5202–eaaw5202 (2019).

41. S. C. Piantadosi, Z. C. Zhou, C. Pizzano, C. E. Pedersen, T. K. Nguyen, S. Thai, G. D. Stuber, M. R. Bruchas, Holographic stimulation of opposing amygdala ensembles bidirectionally modulates valence-specific behavior via mutual inhibition. Neuron 112, 593–610.e5 (2024).

42. J. V. Gill, G. M. Lerman, H. Zhao, B. J. Stetler, D. Rinberg, S. Shoham, Precise Holographic Manipulation of Olfactory Circuits Reveals Coding Features Determining Perceptual Detection. Neuron 108, 382–393.e5 (2020).

43. A. R. Mardinly, I. A. Oldenburg, N. C. Pégard, S. Sridharan, E. H. Lyall, K. Chesnov, S. G. Brohawn, L. Waller, H. Adesnik, Precise multimodal optical control of neural ensemble activity. Nature Neuroscience 21, 881–893 (2018).

44. H. Adesnik, L. Abdeladim, Probing neural codes with two-photon holographic optogenetics. Nature Neuroscience 24, 1356–1366 (2021).

45. A. M. Packer, L. E. Russell, H. W. P. Dalgleish, M. Häusser, Simultaneous all-optical manipulation and recording of neural circuit activity with cellular resolution in vivo. Nature Methods 12, 140–146 (2015).

46. A. Vinograd, A. Nair, J. H. Kim, S. W. Linderman, D. J. Anderson, Causal evidence of a line attractor encoding an affective state. Nature 634, 910–918 (2024).

47. V. Emiliani, A. E. Cohen, K. Deisseroth, M. Häusser, All-Optical Interrogation of Neural Circuits. J Neurosci 35, 13917–13926 (2015).

48. N. N. Foster, J. Barry, L. Korobkova, L. Garcia, L. Gao, M. Becerra, Y. Sherafat, B. Peng, X. Li, J.-H. Choi, L. Gou, B. Zingg, S. Azam, D. Lo, N. Khanjani, B. Zhang, J. Stanis, I. Bowman, K. Cotter, C. Cao, S. Yamashita, A. Tugangui, A. Li, T. Jiang, X. Jia, Z. Feng, S. Aquino, H.-S. Mun, M. Zhu, A. Santarelli, N. L. Benavidez, M. Song, G. Dan, M. Fayzullina, S. Ustrell, T. Boesen, D. L. Johnson, H. Xu, M. S. Bienkowski, X. W. Yang, H. Gong, M. S. Levine, I. Wickersham, Q. Luo, J. D. Hahn, B. K. Lim, L. I. Zhang, C. Cepeda, H. Hintiryan, H.-W. Dong, The mouse cortico–basal ganglia–thalamic network. Nature 598, 188–194 (2021).

49. H. Hintiryan, N. N. Foster, I. Bowman, M. Bay, M. Y. Song, L. Gou, S. Yamashita, M. S. Bienkowski, B. Zingg, M. Zhu, X. W. Yang, J. C. Shih, A. W. Toga, H.-W. Dong, The mouse cortico-striatal projectome. Nature Neuroscience 19, 1100–1114 (2016).

50. J. G. Parker, J. D. Marshall, B. Ahanonu, Y.-W. Wu, T. H. Kim, B. F. Grewe, Y. Zhang, J. Z. Li, J. B. Ding, M. D. Ehlers, M. J. Schnitzer, Diametric neural ensemble dynamics in parkinsonian and dyskinetic states. Nature 557, 177–182 (2018).

51. K. Ade, Y. Wan, M. Chen, B. Gloss, N. Calakos, An Improved BAC Transgenic Fluorescent Reporter Line for Sensitive and Specific Identification of Striatonigral Medium Spiny Neurons. Frontiers in Systems Neuroscience volume 5-2011 (2011).

52. L. Madisen, T. A. Zwingman, S. M. Sunkin, S. W. Oh, H. A. Zariwala, H. Gu, L. L. Ng, R. D. Palmiter, M. J. Hawrylycz, A. R. Jones, E. S. Lein, H. Zeng, A robust and high-throughput Cre reporting and characterization system for the whole mouse brain. Nature Neuroscience 13, 133–140 (2010).

53. S. Gong, M. Doughty, C. R. Harbaugh, A. Cummins, M. E. Hatten, N. Heintz, C. R. Gerfen, Targeting Cre Recombinase to Specific Neuron Populations with Bacterial Artificial Chromosome Constructs. J. Neurosci. 27, 9817 (2007).

54. C. R. Gerfen, D. J. Surmeier, Modulation of Striatal Projection Systems by Dopamine, Annual Review of Neuroscience. 34 (2011)pp. 441–466.

55. M. Maltese, J. R. March, A. G. Bashaw, N. X. Tritsch, Dopamine differentially modulates the size of projection neuron ensembles in the intact and dopamine-depleted striatum. eLife 10, e68041 (2021).

56. M. T. Dang, F. Yokoi, H. H. Yin, D. M. Lovinger, Y. Wang, Y. Li, Disrupted motor learning and long-term synaptic plasticity in mice lacking NMDAR1 in the striatum. Proceedings of the National Academy of Sciences 103, 15254–15259 (2006).

57. A. C. Koralek, X. Jin, J. D. Long II, R. M. Costa, J. M. Carmena, Corticostriatal plasticity is necessary for learning intentional neuroprosthetic skills. Nature 483, 331–335 (2012).

58. X. Jin, R. M. Costa, Start/stop signals emerge in nigrostriatal circuits during sequence learning. Nature 466, 457–462 (2010).

59. F. J. Santos, R. F. Oliveira, X. Jin, R. M. Costa, Corticostriatal dynamics encode the refinement of specific behavioral variability during skill learning. eLife 4, e09423 (2015).

60. J. Dong, S. Hawes, J. Wu, W. Le, H. Cai, Connectivity and Functionality of the Globus Pallidus Externa Under Normal Conditions and Parkinson’s Disease. Frontiers in Neural Circuits Volume 15-2021 (2021).

61. Z. Gu, Z. R. Lewis, J. Tang, A. Mendelsohn, A. M. Vicente, L. Nikoobakht, S. Rosenberg, A. Chakravarth, V. Paixao, M. S. Tirumalasetti, G. A. Abuagala, A. Khalili, E. D. Thomas, K. A. Fancher, M. Mallory, A. Manning, D. Bertagnolli, J. Goldy, C. Rimorin, M. Tieu, T. Cardenas, A. Torkelson, A. B. Chakka, S. Yao, S. Sorensen, K. A. Smith, L. A. Hammond, D. S. Peterka, H. Zeng, B. Tasic, R. M. Costa, The External Globus Pallidus is a Basal Ganglia Output Hub with Action-Specific Circuits. [Preprint] (2026). 10.64898/2026.04.12.718022.

62. H. H. Yin, S. P. Mulcare, M. R. F. Hilário, E. Clouse, T. Holloway, M. I. Davis, A. C. Hansson, D. M. Lovinger, R. M. Costa, Dynamic reorganization of striatal circuits during the acquisition and consolidation of a skill. Nature Neuroscience 12, 333–341 (2009).

63. S. Arber, R. M. Costa, Networking brainstem and basal ganglia circuits for movement. Nature Reviews Neuroscience 23, 342–360 (2022).

64. P. Redgrave, T. J. Prescott, K. Gurney, The basal ganglia: a vertebrate solution to the selection problem? Neuroscience 89, 1009–1023 (1999).

65. S. Grillner, B. Robertson, M. Stephenson-Jones, The evolutionary origin of the vertebrate basal ganglia and its role in action selection. The Journal of Physiology 591, 5425–5431 (2013).

66. A. Falasconi, H. Kanodia, S. Arber, Dynamic basal ganglia output signals license and suppress forelimb movements. Nature 644, 749–758 (2025).

67. V. R. Athalye, J. M. Carmena, R. M. Costa, Neural reinforcement: re-entering and refining neural dynamics leading to desirable outcomes. Current Opinion in Neurobiology 60, 145–154 (2020).

68. R. M. Neely, A. C. Koralek, V. R. Athalye, R. M. Costa, J. M. Carmena, Volitional Modulation of Primary Visual Cortex Activity Requires the Basal Ganglia. Neuron 97, 1356–1368.e4 (2018).

69. V. R. Athalye, K. Ganguly, R. M. Costa, J. M. Carmena, Emergence of Coordinated Neural Dynamics Underlies Neuroprosthetic Learning and Skillful Control. Neuron 93, 955–970.e5 (2017).

70. V. R. Athalye, P. Khanna, S. Gowda, A. L. Orsborn, R. M. Costa, J. M. Carmena, Invariant neural dynamics drive commands to control different movements. Current Biology 33, 2962–2976.e15 (2023).

71. V. R. Athalye, F. J. Santos, J. M. Carmena, R. M. Costa, Evidence for a neural law of effect. Science 359, 1024–1029 (2018).

72. T.-C. Kao, G. Hennequin, Neuroscience out of control: control-theoretic perspectives on neural circuit dynamics. Current Opinion in Neurobiology 58, 122–129 (2019).

