## Supplemental Information for "Striatal ensembles specify and control ongoing actions at fine motor resolution"

Science
AAAS

Supplementary Materials for

### Striatal ensembles specify and control ongoing actions at fine motor resolution

Ines Rodrigues-Vaz, Vivek R. Athalye, Helio F.M. Rodrigues, Darcy S. Peterka, Rui M. Costa

**The PDF file includes:**

Materials and Methods
Supplementary Text
Figs. S1 to S4
Tables S1 to S3
References

#### Materials and Methods

Animals

All experiments and procedures were performed according to National Institutes of Health (NIH) guidelines and approved by the Institutional Animal Care and Use Committee (IACUC) of Columbia University. Adult mice, aged 2–6 months, were used for these experiments. In order to label D1- and D2-spiny projection neurons (SPNs, which are also termed medium spiny neurons, MSNs), we used the following transgenic lines (table S3): Tg(Drd1-cre)EY217Gsat (RRID:MGI:4366805)(*53*), Tg(Drd1-cre)EY217Gsat crossed with Ai9(RCL-tdT) (RRID:IMSR_JAX:007905)(*52*), Tg(Adora2a-cre)KG139Gsat (RRID:MMRRC_036158-UCD)(*53*), Tg(Adora2a-cre)KG139Gsat crossed with Ai9(RCL-tdT) and Drd1a-tdTomato line 6 (RRID:IMSR_JAX:007905)(*51*). For experiments testing the necessity of plasticity in the striatum we used the following lines (table S3): RGS9-cre (RRID:IMSR_JAX:020550)(*56*) crossed with NMDAR1-loxP (RRID:IMSR_JAX:036352)(*56*). For comparison, we used the negative Cre littermates as controls.
Mice used for experiments were individually housed and kept under a 12-h light-dark cycle.

Randomization and blinding

The experimenter was blinded to the experimental groups for plasticity experiments.
All animals were run on each day in a fixed random order.
Holographic manipulations were performed according to a pre-defined protocol and controlled by Matlab in closed-loop with the mouse's behavior - not manually by the experimenter.
Sessions for stimulating D1- and D2-SPNs were interleaved as much as possible.

Stereotaxic Surgery

Surgical procedures for imaging of neurons in the dorsolateral striatum (DLS) were performed in two steps consisting of viral injection at least 5 days prior to lens implant, following protocols described in protocols.io ("Viral injection" and "Lens implant").
Before each surgery, analgesia was administered to ensure 72 hours of pain relief. The mouse was placed in a stereotaxic frame under isoflurane anesthesia on a heated pad.
For viral injection, capillary tubes (Drummond 3.5" #3-000-203-G/X) were pulled using a Pipette puller (Sutter P-2000; Heat 360, FIL 5, Vel 25, Del 10, Pul 50) and backfilled with mineral oil. A craniotomy was made using a dental drill, and the capillary tube was lowered to the target coordinates. For imaging, 300 nl of AAV5.CaMKII.GCaMP6f (titer: $1\times10^{12}$–$1\times10^{13}$ GC/ml) was injected into the left DLS (AP: +0.75 mm, ML: 2.5 mm, DV: 2.4 mm) at a rate of ~5 nl/pulse. For holographic optogenetics experiments, an additional 300 nl of AAV8.CaMKII.ChRmine (titer: $\sim1\times10^{11}$ GC/ml) was co-injected with AAV5.CaMKII.GCaMP6f at a 1:1 ratio into the left DLS at the same coordinates. Following injection, the capillary tube was left in place for at least 5 minutes before slow withdrawal, and the skin was sealed with sutures.
At least 5 days after viral injection, a 1 mm GRIN lens implant surgery was performed. Following craniotomy and cortex removal – using vacuum aspiration with 1 mm blunt needle – the GRIN lens was implanted at the same coordinates as the viral injection at depth of DV: 2.0 mm. A head bar was secured and a protective cap was built around the lens, which was optionally covered with silicone sealant (Kwick-Sil or Kwick-Cast) for protection.
For experiments requiring electromyographic (EMG) recordings of forelimb muscle activity, we fabricated EMG electrodes in-house as previously described (*72*) and in protocols.io ("EMG

fabrication”). Briefly, two pieces of insulated braided stainless-steel wire were cut, knotted together, and secured to a working surface. Under a microscope, 0.5 mm portions of insulation were stripped from each strand just below the knot, leaving exposed contact sites separated by 0.5 mm. The wire ends were inserted into hypodermic needles and crimped in place to facilitate muscle insertion, and the opposing ends were soldered into corresponding sites on a miniature Omnetics connector.

EMG electrodes were then implanted into forelimb muscles (biceps, triceps, palmaris longus, and extensor digitorum communis) following lens implant (*42*), as described in protocols.io ("EMG implant"). Prior to surgery, mice were habituated to an Elizabethan collar and gel food for 4 days to minimize post-surgical disruption. On the day of surgery, analgesia was administered as described above. Electrodes were inserted into each target muscle using the hypodermic needle as a guide and secured in place by knotting the wire at the exit point to fix the recording sites within the muscle. Each incision was closed with sutures and covered with antibiotic ointment. The EMG connector was attached to the back of the skull or existing head cap using dental cement (Metabond), and any neck incision was similarly closed with sutures and antibiotic ointment. After surgery, the Elizabethan collar was placed on the mouse for at least 5 days. Food gel and soften pellets were provided for at least 5 days post-surgery.

<u>Histology</u>

Histology was performed as described in protocols.io (Histology) to verify lens coordinates and viral expression. Cardiac perfusion used 4% PFA solution to ensure fixation of the brain tissue. At least 24 hours after the cardiac perfusion, the brain was moved into 0.05% Thimerosal-PBS and stored in the fridge until histological processing.

Brains were sliced coronally at 50 µm using a vibratome (Leica VT1000) and collected into 1X PBS. Immunohistochemistry was performed as follows:

- To localize the imaging site, sections were incubated with an Alexa Fluor 488-conjugated GFP antibody at 1:1000 in 0.4% Triton-PBS overnight at room temperature.
- To identify tdTomato somatic expression, sections were incubated with a primary anti-RFP antibody at 1:2000 in 0.4% Triton-PBS overnight at 4°C, followed by a secondary anti-Rabbit Alexa Fluor 647 antibody at 1:1000 in 0.4% Triton-PBS for 2 hours at room temperature.

All sections were counterstained with DAPI at 1:1000 in PBS for 15 minutes and then mounted on glass slides with Mowiol solution, allowed to dry overnight, and sealed with nail polish. Sections were imaged using a custom-built automated slide scanner equipped with a 4x 0.4-NA Plan Apo objective (Nikon Instruments Inc.) and P200 slide loader (Prior Scientific), controlled by NIS-Elements using custom acquisition scripts, from the Zuckerman Institute's Cellular Imaging Platform. For detection of DAPI, Alexa Fluor 488, and Alexa Fluor 647, the following excitation wavelengths and filters were used, respectively: 405 nm (filter 435/26 nm), 488 nm (filter 525/30 nm), and 647 nm (filter 705/72 nm). Image processing and analysis were performed using BrainJ (*73*) (RRID:SCR_027061).

<u>Two-action isometric task</u>

<u>*Setup*</u>

Mice were head-fixed and placed in a 3D-printed cup (*74*) with a sucrose spout placed within tongue reach. The right forelimb interacted with an immobile joystick (3 mm screw) mounted on 3D-printed frames attached to two load cell sensors (Phidgets 3139_0), one measuring push/pull and the other measuring left/right forces. The left forelimb rested on an auxiliary pole equipped with two additional load sensors measuring push/pull and vertical forces. The cup holding the mouse was mounted on a fifth load sensor to capture overall body movement.

Signals from all load sensors were recorded at 1 kHz using the Champalimaud Foundation Scientific (CFS) "Scientific Board v1.3" with "Quad Load Cell v1.2" module, which also detected threshold crossings based on experimenter-defined parameters. Data were visualized and stored using the CFS "Load Cells Visualizer" software.

A pyControl "Lickometer v1.0" board detected licking and contact with the joystick and auxiliary pole. Task control and sucrose delivery were managed by a pyControl (*75*) "Breakout 1.2" board, which logged events at 1 kHz and opened solenoid to dispense the reinforcer - 5 µl of 10% sucrose solution. Sucrose flowed via gravity through tubing and was delivered through a blunt 16G needle spout positioned near the animal's mouth. Solenoid timing was calibrated before each session, and the sucrose solution was prepared fresh weekly. Hardware schematics are shown in Fig. S4a.

*Training*

Mice were habituated to head fixation for 15 min on a running wheel over four days and then food-restricted to 80–85% of initial body weight. Training for the two-action isometric task was performed as described in protocols.io ("Behavior training"). Mice pushed or pulled an immobile, pressure-sensitive joystick in a self-paced manner to receive reinforcer. A task action was defined as a force exceeding a task-defined threshold for longer than the task-defined duration. Thresholds, durations, and reinforcement delays were progressively adjusted during training – these are referred to as "Task Parameters" (table S4). Reinforcement followed the end of a valid action after a delay. A 3 g quiescence threshold was required for trial initiation.

Each trial consisted of: (1) a 1 s inter-trial interval (ITI); if forces were below 3 g at 1 s, a new trial began, otherwise the ITI reset; (2) a trial period, during which a task action (push or pull exceeding the threshold for the task duration) triggered delayed sucrose delivery. Trials terminated upon reinforcement or when force fell below task-defined threshold after an action that did not meet task-defined duration.

Training comprised three sequential blocks with distinct reinforcement contingencies. In Block Both, push and pull actions were both reinforced, up to 50 reinforcements per action per session. On the final session of Block Both, the less frequent action was designated action A and the more frequent as action B. In Block A, only action A was reinforced – starting after last session of Block Both; in Block B, only action B was reinforced – starting after last session of Block A. Block Both sessions ended after 50 reinforcements of each action or 30 min. Blocks A and B ended after 200 reinforced actions or 30 min. Task parameters were adjusted during training, advancing according to table S4 when mice executed >50 reinforced task actions in a session. Block Both concluded when mice exceeded 50 reinforced actions in two consecutive sessions at Parameters 3. Blocks A and B each concluded when the reinforced action comprised ≥80% of total actions - defined as reinforced task actions divided by total action A and action B executions - over three consecutive sessions at Parameters 3. Conclusion of Block B marked the end of training, after which animals were returned to ad libitum feeding.

For Rgs9-LCre::Grin1tm1Yql homozygous mice (Striatal NR1-KO) and Cre-negative control littermates, the same training protocol was used, with two modifications. Each block was capped at 1.2× the number of sessions required by the third fastest mouse in the cohort to reach Parameters 3 (table S4). Animals were also advanced to the next block if they met give-up criteria, defined as fewer than ten 3g threshold crossings (≥50 ms duration) averaged over three consecutive sessions.

Selected sessions for analysis of behavioral training

Because animals were trained to criterion, the number of sessions per block varied across individuals. To enable cross-animal comparisons, specific sessions were selected for analysis. In Block Both, four sessions were chosen, and in Blocks A and B, five sessions were chosen, following the selection rules detailed in table S5. These sessions were used in Fig. S2 and S3.

Adaptive Isometric task

*Setup*

Mice were trained on this task to test causal striatal control of specific ongoing actions. To do that, we developed a modified version of the two-action forelimb task that promoted balanced performance of both actions. Reinforcement probability was adjusted online based on recent performance: the proportion of push and pull actions was tracked over the last 10 trials. When both actions occurred in fewer than 70% of recent trials, both were reinforced; if one action exceeded 70%, that action was unrewarded.

The setup was otherwise identical to the two-action isometric task (Fig. S4A), with additional hardware for closed-loop holographic stimulation (Fig. S4B). This system required flexibility to: (1) select, on a single-trial basis, the threshold crossing to trigger stimulation; (2) trigger laser with millisecond latency; and (3) specify the holographic 2-photon pattern, on a single-trial basis. Features 1 and 2 required an improved hardware system for processing force sensors, while feature 3 required a software update to the two-photon microscope's stimulation interface. Features 1 and 2 were accomplished using an updated load-cell system ("HARP Load Cells Acquisition v1.1" and "HARP Load Cell Interface v1.1," CFS Hardware Platform), which evaluated in parallel when the force exerted on the joystick crossed six experimenter-defined thresholds. In the holographic experiments, six thresholds were defined: 3 g, 4 g, and 6 g for push and pull actions. To select which threshold would trigger stimulation on a given trial, and to restrict the trigger signal to specific time windows within each trial, a custom circuit board consisting of a MUX and an AND gate was built to route the appropriate trigger signal to the microscope while minimizing the delay between threshold crossing and stimulation onset. The MUX and AND gate was controlled by custom MATLAB scripts running on the microscope control computer, which determined which force threshold should serve as the trigger on each trial and when to enable it. Digital communication between the task boards, the computer, and the MUX and AND gate was implemented using a NIDAQ DIO board (USB-6001).

Feature 3 was accomplished through a newly implemented function in the PrairieLink API of the Bruker microscope, which allowed two-photon stimulation patterns to be specified dynamically during an ongoing imaging experiment. Schematics of the full setup, including both the behavioral and stimulation hardware, are presented in Fig. S4.

*Training*

This task was used for holographic stimulation experiments and followed the two-action forelimb task with modified reinforcement to promote balanced performance of both actions. Mice were trained for 3 weeks with adaptive reinforcement and no stimulation. Training was conducted partly in behavior boxes in the absence of imaging, prior to head-fixed sessions under the microscope.

2-photon microscope

The optical setup comprised two femtosecond pulse lasers coupled to a custom-modified two-photon laser scanning microscope (Ultima In Vivo, Bruker Corporation, Billerica, Massachusetts) equipped with an 8 kHz resonant scanner. The laser source for imaging was a pulsed Ti:sapphire laser (Chameleon Vision-S, Coherent, Inc., Saxonburg, Pennsylvania) tuned to 920 nm for GCaMP6f imaging, with power controlled by a Pockels cell (350–105-BK, 302RM controller, Conoptics, Inc., Danbury, Connecticut). The laser beam was expanded by a 1:2 telescope (Thorlabs GBE02-B) and further scaled in the microscope by a 1:1.33 telescope before being coupled into a scan lens (f = 75 mm), tube lens (f = 180 mm), and objective lens (16X 0.8NA N16XLWD-PF, Nikon Corporation, Tokyo, Japan). The laser could also be directed to a non-resonant scanning path where both X and Y scanning were controlled by galvanometric mirrors. The fluorescence signal was collected through the objective lens, split from the IR beam with a 670 nm long-pass dichroic (ZT670rdc, Chroma Technology Corp., Bellows Falls, Vermont), and coupled into a liquid light guide assembly. The light was split by a 565 nm long-pass dichroic mirror (HQ565dcxr, Chroma Technology Corp.), with the green and red channels each directed to GaAsP PMTs through 510/20 and 607/45 bandpass filters, respectively (Chroma Technology Corp.).

The photostimulation path was largely independent from the imaging path. The laser source for photostimulation was a low repetition rate (1 MHz) amplified laser (Monaco 1035–40, Coherent, Inc., Saxonburg, Pennsylvania) operating at 1035 nm, with power controlled by an integrated acousto-optic modulator. The beam was expanded by a 1:2 telescope (Thorlabs GBE02-B) and directed into the microscope, where it could be steered either by a dedicated pair of galvanometers or coupled to a customized light path including a spatial light modulator (Bruker Neurolight 3D, Billerica, Massachusetts) for holographic stimulation. The photostimulation beam was combined and made coaxial with the imaging beam just before the scan lens via a 1030 nm short-pass dichroic (T1030SP, Chroma Technology Corp.).

Imaging and photostimulation were controlled by PrairieView (Bruker Corporation, Billerica, Massachusetts). Custom software defined stimulation patterns and triggered closed-loop photostimulation based on behavioral events - MATLAB (The Mathworks, Inc., Natick, Massachusetts) and Python, interfaced with PrairieView via the PrairieLink API.

2-photon imaging

Before each experimental cohort, the imaging and stimulation laser powers were measured at the output of 16X Nikon objective and confirmed stable across months. A custom computerized goniometer was used to precisely and reproducibly angle the head of the mouse such that the imaging lens was orthogonal to the beam path.

2-photon imaging was performed as described in protocols.io ("2-photon imaging"). Prior to each session, laser alignment was confirmed at the objective output. The mouse was head-fixed in the behavioral setup under the microscope, and the top of the GRIN lens was located and cleaned if necessary. Imaging power was continuously monitored using a beam pickoff (Thorlabs BSF10-B) and power meter (Thorlabs PDA100A2), with the Pockels cell bias adjusted accordingly throughout the session.

The typical imaging power was <50 mW, and could be up to 80 mW for imaging deeper than ~250 µm below the terminus of the GRIN lens. Images were acquired at 30 Hz over a 512 × 512 pixel field of view. For Fig. 1 and 2, a 1.5x zoom was used over 496.1 µm × 496.1 µm; for Fig. 3

and 4, a 1.5x zoom over 550.4 μm × 550.4 μm and a 2x zoom over 412.7 μm × 412.7 μm were used.
For cell type identification, 1000 frames of functional and structural images were acquired using green and red PMTs respectively. These data were then processed as described below (Methods – Distinguishing D1- and D2-SPNs). Structural images and z-stacks were acquired at the end of each session using both green and red channels to confirm ROI identity (Fig. S1).
Most mice were imaged for the entire duration of training, with the same field of view tracked across sessions. ROIs identified in previous sessions were used to help locate the same field of view, and custom code was written to track the same ROIs across days.

<u>Protocol to image the same field of view and neurons across sessions</u>
For holographic stimulation experiments, the same field of view and neurons were imaged across paired sessions. After the first session, imaging data were processed with Suite2p (*76*) to extract active ROIs. In the paired session, PrairieView's "Brightness over Time" (BOT) feature was used to annotate these neurons in the GUI. XYZ coordinates were adjusted to best match the previous session's field of view.

<u>Calibration of 2-photon stimulation spatial targeting and power</u>
When doing holographic stimulation, the spatial targeting of 2-photon stimulation was calibrated before each session (protocols.io, "2-photon stimulation calibration"). Using PrairieView's "burn spots" routine, we iteratively aligned burned spots on a fluorescent sample to target locations, allowing the software to calibrate galvos' control for accurate stimulation. Calibration was stable across sessions, requiring only minor adjustments. This calibration was done using the same objective used for the experimental recordings.
Before experiments, we calibrated stimulation power and duration for each ROI. Using custom Python scripts with PrairieView's "BOT" feature, we visualized ROI responses while sweeping power (3–7 mW per target) and duration (50-100 ms) to identify the minimum parameters that reliably activated the targeted neurons.

<u>Holographic optogenetics experiments</u>
We performed holographic optogenetics experiments to test the effect of stimulating specific neuronal ensembles on self-paced forelimb actions during the Adaptive Isometric task. Experiments relied on pairs of sessions in which we imaged the same field of view and neurons. On the first session, we performed two-photon imaging as mice performed the task, used Suite2p to extract functional ROIs, and automatically labelled ROIs as D1- or D2-SPNs. On the second session, we performed the stimulation experiment, consisting of a calibration block followed by a stimulation block (Fig. 3F). For each stimulation session, cell type choice (D1- or D2-SPNs) was determined by balancing the number of sessions per cell type and the number of neurons available for stimulation.
Within the calibration block, we performed two-photon imaging as mice performed the adaptive isometric task for up to 30 minutes. Using PrairieView's "BOT" feature, we pre-defined ROIs identified from the previous session and rapidly extracted their fluorescence, averaged over the pixels within a circle defined by each Suite2p mask's center and radius. We extracted behavioral data and identified force threshold crossings of 3 g (hereafter "3g cross actions"). For either D1- or D2-SPNs, we z-scored each ROI's fluorescence across the recording and fit an SVM to predict push versus pull using single trials of neural activity averaged over the 5 imaging frames

centered at force peak. If SVM accuracy on test trials exceeded 55%, we used the SVM weights to identify push- and pull-weighted ensembles with equal numbers of neurons, and plotted their activity time-locked to the force peak of 3g cross actions. If the push (pull) ensemble showed greater activity for push (pull) actions, we proceeded to holographic stimulation.
We then defined a software command via the PrairieLink API to specify holographic stimulation for each ensemble. Stimulation was targeted to the center of each ROI in the ensemble, with total power calculated as the product of the number of neurons per ensemble and the desired average power per neuron. Stimulation duration was typically 100 ms - sufficient to reliably activate neurons expressing ChRmine. To cover each ROI's circular extent, dedicated galvos traced 5 spirals per stimulation with a diameter matching the average ROI diameter. Stimulation was configured to be delivered upon a trigger signal corresponding to a force threshold crossing.
Within the stimulation block, we performed two-photon imaging and closed-loop holographic stimulation as mice performed the adaptive isometric task for 30 minutes. We pre-defined a schedule interleaving conditions based on which action triggered stimulation (3g cross push or pull) and which stimulation pattern was delivered (push ensemble, pull ensemble, or no stimulation). Trials were organized into alternating blocks of 6, triggered by one action (either 3g cross push or 3g cross pull), with the following stimulation sequence per block: (1) no stimulation, (2) push ensemble, (3) no stimulation, (4) no stimulation, (5) pull ensemble, (6) no stimulation. A 2-second inter-trial interval was imposed between trials.
Our custom setup (Fig. 3 and S4B) enabled selection of which force threshold crossing served as the trigger on interleaved trials and stimulation delivery within milliseconds of threshold crossing — critical given the rapid timescale of force changes. As described in the setup section above, the microscope computer ran custom MATLAB code to step through the scheduled trials, using the PrairieLink API to send stimulation commands and a NIDAQ DIO board (USB-6001) to select and enable force thresholds via the MUX and AND gate. The trigger signal output from the MUX was also received by the computer to track stimulation delivery and advance to the next inter-trial interval and trial.

EMG recordings

EMG recordings were performed as described in protocols.io ("EMG recordings"). Custom-made electrodes were implanted in forelimb muscles and connected to an Omnetics connector. Prior to each session, mice were head-fixed in the behavioral setup under the microscope and the 16-channel ZC16 headstage (Tucker-Davis Technologies – TDT) was connected to the implanted connector. EMG recordings were set to true differential mode in Synapse recording software, and the signal quality was confirmed when the mouse moved. Signal quality was optimized by grounding the mouse to the air table. Signals were acquired via the ZC16 headstage connected to a TDT RZ5D Bioamplifier and PZ5-32 preamplifier, and recorded at 24 kHz. For analysis, EMG signals were downsampled to 1 kHz, high-pass filtered at 40 Hz, rectified, and smoothed with a Gaussian kernel (SD = 25 ms) (*77*). Data included 41 recording sessions from 4 mice performing the two-action isometric task (Fig. 1).

Quantification and Statistical Analysis

Analysis was performed using custom scripts in Python.

Data extraction and synchronization

All data streams—task events (1 kHz), joystick forces (1 kHz), EMG (24 kHz), 2-photon imaging/voltage (30 Hz/1 kHz) - were synchronized via the ITI signal, providing a unified timestamp across modalities for precise alignment of behavior, forces, EMG and imaging.

Calibrated force measurement

The joystick voltage signal was converted to force using a linear model:

Force (g) = (Voltage – baseline) × conversion factor

Baseline voltage was measured before each session and defined as the average voltage with no weight on the sensors. The conversion factor was determined by applying known weights (0, 2, 5, 20 g) and fitting a linear model. Conversion was stable and re-calibrated weekly to correct drift.

Definition of actions

For analysis, we defined several actions from force and touch signals. Task actions were push and pull events identified by the hardware from 1 kHz force traces, with intervals from threshold rise to fall. 3 g cross actions were pushes or pulls exceeding 3 g for ≥66 ms with joystick touch ≥100 ms before crossing to ensure isometric behavior; force traces were downsampled to 30 Hz. 6 g cross actions were a subset of 3 g crosses peaking above 6 g, the final training threshold. Touch was defined as joystick contact <2 g for >1 s, and licks as ≥5 licks with <0.5 s inter-lick interval, excluding overlap with 3 g crosses. We also analyzed special subsets: pre-isolated 3 g crosses had no other 3 g crosses in the 0.5 s before, post-isolated 3 g crosses had none in the 0.5 s after, and matched actions were push/pull trials matched for continuous-valued features (Methods - Matching trials across actions).

Action events

For push and pull actions, analysis was aligned to two key events within the action interval: the moment the force threshold was crossed (rising edge) and the moment of peak force. For touch and lick actions, analysis was aligned to the midpoint of the action interval.

Matching trials across actions

We tested whether muscle and neural activity encoded push and pull actions using trials matched for average features (Fig. 1F-H and S3). Tolerances were: average force ≤0.1 g (Fig. 1F-H); peak force ≤0.1 g, two-axis force magnitude at peak ≤0.2 g, action duration ≤30 ms, and lick rate/probability during actions and at peak ≤0.5 Hz and ≤0.1, respectively (Fig. S3).
Trials were iteratively removed to achieve matching while minimizing loss of trials. Features were z-scored, then for each iteration: (1) define action 1 as the one with fewer trials, action 2 as the other; (2) compute the target vector as the difference in mean features across actions; (3) zero in-tolerance features (i.e. set the difference to zero for each feature that is within-tolerance); (4) compute dot products of action 2 trials with the target; (5) remove the trial with the smallest dot product; (6) repeat until all features met tolerance.
For Fig. S3, trials were pre-selected such that the time between solenoid opening and force threshold crossing was at least 4 seconds, ensuring that analyzed actions did not occur during sucrose consumption**.**

Time course of action force

Trial-averaged joystick force was computed as peri-event time histograms (PETHs) locked to action events: 6 g cross push/pull actions were aligned to force peak (Fig. 1C). In Fig. 2F, -4–0 s shows pre-isolated and 0–4 s shows post-isolated 3 g cross force, illustrating trials used to predict action identity from neural activity.

Quantification of actions across reinforcement

Action performance was quantified across the reinforcement schedule. For task actions, we quantified action rate for each session (Fig. S2B,C). For 6 g cross actions, we quantified action rate pooled over sessions in each reinforcement block (Fig. 1D). For trials matched across 3g cross actions, we quantified number of actions, peak and average force, two-dimensional force magnitude, average lick probability, action duration, action rate, and action proportion (Fig. S3A-H).

For Striatal NR1-KO mice, we quantified 6g cross actions with the overall action rate for each session (Fig. S2H), and the action rate pooled over sessions in each reinforcement block (Fig. S2I).

EMG analysis

We analyzed pre-processed EMG (Methods – EMG Recordings) from four forelimb muscles for 6 g cross actions with force matched within 0.1 g. EMG channels were z-scored across each session and aligned to the 6 g cross event. We compared the vector magnitude of EMG difference across actions to the difference within each individual action. Concretely, for each session, we split trials for each action into two halves and calculated the EMG PETH for each half. At each time point locked to 6g cross, we calculated the vector magnitude of difference across halves within an action, as well as the difference across actions using the PETH from one half. We compared across-action and within-action difference in the 100ms centered at 6g cross (Fig. 1H).

Neuron detection and activity extraction with Suite2p

2-photon imaging data were processed with Suite2p (*76*) for motion correction, neuron detection, and fluorescence extraction; ROIs were classified using a classifier trained on sparse striatal SPN activity, neuropil-corrected ($F_{corrected} = F – 0.7 \times F_{neu}$), and neural activity was reported as z-scored $\Delta F/F_0$ with $F_0$ as the 1-minute running median. Fig. 3H,I,K-M shows z-scored, raw fluorescence averaged over ROI pixels from PrairieView's "BOT".

Distinguishing D1- and D2-SPNs

We developed an automated thresholding method to identify D1- and D2-SPNs based on fluorophore expression (Fig. S1). Functional imaging (green channel) was used to detect ROIs with Suite2p, while structural imaging (red channel) determined co-localization of cell-type–specific fluorophores. The overall workflow is outlined in Fig. S1A. The method involved three main steps: (1) aligning and processing structural and functional data, (2) labeling ROIs manually (ground truth) and automatically using red-channel thresholds, and (3) comparing manual and automatic labeling.

1) Structural data processing:
   Functional data were motion-corrected and ROIs extracted using Suite2p (Fig. S1B). As fluorophore expression is soma-restricted, only pixels within a 12-pixel diameter circle of each ROI center were used. Structural images (red channel, 1000 frames) were motion-

corrected against the functional template, then background-subtracted using a 25-pixel rolling ball filter in ImageJ (Fig. S1C,D).

2) ROI labeling:
ROIs near the edge of the GRIN lens were excluded to avoid aberrations; the center of the lens was estimated from the red channel by computing the median coordinates of pixels above the 20th percentile intensity. Valid ROIs fell within a circle around this center.
   a. *Manual labeling:*
   Manual labels were assigned on average, motion-corrected, background-subtracted structural images overlaid with functional ROIs (Fig. S1E). Labels were “red,” “non-red,” or “no-ID” if uncertain. Each mouse had at least two sessions labeled three times (folds); only ROIs consistently labeled across all folds were retained (Fig. S1F,G). Thresholds of 0.7 (red) and 0.3 (non-red) were used for comparisons in Fig. S1.
   b. *Automatic labeling:*
   Structural images were processed with Ilastik (*78*) (v1.3.3) to estimate per-pixel probability of being red (Fig. S1H). ROIs with >0.7 red pixels were labeled “red,” <0.3 as “non-red,” and 0.3–0.7 as “no-ID” (Fig. S1I).

**3)** Manual vs. automatic comparison:
Label accuracy, swapping errors, and single-method errors were quantified across threshold pairs to optimize performance (Fig. S1J-M).

Thresholds of 0.3/0.7 (non-red/red) were used for general characterization (Fig. 2), and 0.45/0.55 thresholds were used for holographic stimulation experiments (Fig. 3 and 4).

Time course of neuronal activity

Neuronal activity was analyzed time-locked to action events. An example trace of activity averaged over D1-, D2-, and all SPNs is shown alongside force (Fig. 2B). We analyzed session-averaged PETHs aligned to the force peak of pre-isolated 3 g cross actions (Fig. 2C). We specifically analyzed activity at force peak and 0.5 seconds before force peak, computed as z-scored F/F0 averaged over 3 frames centered at each time point. Averaging over neurons of each cell type, we compared activity at force peak versus 0.5 seconds before force peak (paired t-test). We also compared population-averaged activity across actions at force peak (paired t-test). We plotted PETHs of example individual D1- and D2-SPNs (Fig. 2D). Only sessions with ≥20 trials per action and ≥10 neurons were analyzed (same as in Methods – Support Vector Machine (SVM) decoder model).

Support Vector Machine (SVM) decoder model

To test whether SPNs encoded specific actions, we trained a linear SVM (sklearn.svm.LinearSVC) to predict action identity (e.g. push versus pull) from single-trial population activity. Each trial vector comprised the $\Delta F/F_0$ of each neuron averaged over 5 frames (~167 ms) centered on the action event (i.e. the frame at the event and 2 frames before and after) (Fig. 2H,I and S2). Only sessions with ≥20 trials per action and ≥10 neurons were analyzed, and trials were weighted by 1/(number of trials per action class) to prevent bias. Trials were randomly split 90/10 for training/testing, repeated 100 times. The regularization parameter (C) was optimized over a range of $1\times10^{-5}$ to 100 via 10-fold cross-validation on the training set (with 80% of training data used for sub-training and 20% for sub-testing in each sub-fold). Accuracy was computed for each action separately and overall, and compared to a shuffle control in which the model was trained on randomized action labels but tested on non-randomized labels**.**

Matching number of D1- and D2-SPNs for SVM decoding

To compare D1- and D2-SPNs, we equalized their numbers for SVM decoding (Fig. S3K). For the cell type with more neurons, a random subset was selected in each training/testing fold to match the number of neurons in the other cell type.

SVM decoding across sessions

We reported SVM decoding accuracy for individual sessions and pooled across sessions, under different reinforcement conditions.

*SVM accuracy across each pair of actions, pooling the selected sessions across all reinforcement blocks:*

- Fig. 2H decoded the population activity vector including all SPNs. For push and pull, 3g cross actions were used, and neural activity was centered at force peak.
- Fig. 2I is the same as Fig. 2H, but decoding population activity of a cell type (D1- or D2-SPNs).
- Fig. S3I is the same as Fig. 2H, and Fig. S3J is the same as Fig. 2I, except for matched 3g cross actions and separating by action A and B rather than push and pull.

*SVM accuracy for each selected session across reinforcement:*

- Fig. S2D decoded all SPNs at force peak for 3g cross actions, predicting push versus pull.
- Fig. S2E (left) is the same as Fig. S2D, but decoding population activity of a cell type.
- Fig. S2F (left) is the same as Fig. S2D, except with one SVM predicting action A versus touch and another SVM predicting action B versus touch.

*SVM accuracy pooling selected sessions within each reinforcement block:*

- Fig. S2E (right) decoded population activity of a cell type at force peak for 3g cross actions, predicting push versus pull.
- Fig. S2F (right) decoded all SPNs at force for 3g cross actions, with one SVM predicting action A versus touch and another SVM predicting action B versus touch.
- Fig. S2G is the same as Fig. S2F (right), but decoding population activity of a cell type.
- Fig. S2J is the same as Fig. S2E (right), except decoding all SPNs for NR1-KO and littermate control mice.

Time course of action decoding

We used the SVM to analyze the time course of action decoding. We extracted and decoded neural activity centered at specific time points relative to action events. In Fig. 2E-G, we analyzed non-overlapping 5 frame bins locked to force peak. The bin at time 0 was centered at force peak. From -4 seconds to 0 seconds, the SVM was applied on trials from pre-isolated 3g cross actions. From after 0 seconds to 4 seconds, the SVM was applied on trials from post-isolated 3g cross actions. The SVM used all SPNs in Fig. 2EF and matched numbers of D1- or D2-SPNs in Fig. 2G. In table S1, ref. 2.45-2.53, we analyzed 5 frame bins with a temporal step of 2 frames (~67ms) locked to 3g cross of pre-isolated 3g cross actions.

Overlay of decoder accuracy and force

Decoder accuracy and force were visualized in normalized units (Fig. 2F) by centering data at -4 seconds from force peak (i.e. subtracting the force at -4 seconds from all time points) and scaling by the window's (-4 s to +4 s) maximum trial-averaged value (i.e. dividing by the maximum).

SVM weights and dimension

The SVM assigned positive (negative) weights to neurons whose activation biased the SVM to predict push (pull). For analysis of the weights directly, we z-scored the weights for each decoder (Fig. 3G,J). The weights define a dimension of population activity that best discriminates action identity. We extract this dimension by using the decoder's weight vector and dividing by its L2 norm. This is a vector $w \in R^N$, where $N$ is the number of neurons and the vector magnitude is 1 ($\|w\|_2 = 1$).

Time course of SVM projection

We visualized the time course of action-specific population activity by projecting neural activity onto the SVM dimension. At each time point, the vector of population activity $x \in R^N$ (where each entry is one neuron's activity) was projected onto the SVM dimension: $w^T x \in R^1$, resulting in a scalar that represents the weighted sum of neuronal activity. In Fig. 2J, we used the SVM trained at force peak and projected activity from pre-isolated 3g cross actions time locked to force peak. In Fig. 3H,I and N, we used the SVM trained at force peak and projected activity locked to holographic stimulation.

SVM-weighted neural ensembles

We defined neuronal ensembles that biased the SVM toward push or pull. For each action, we selected the top $k *$ ensemble_frac neurons with the largest positive (push) or negative (pull) weights, where $k$ is the smaller of the number of push- or pull-weighted neurons. We used ensemble_frac $= 0.25$ to visualize the most strongly weighted neurons (Fig. 2K-N) and ensemble_frac $= 1$ to select as many neurons as possible for holographic optogenetic stimulation (Fig. 3G-N).

Time course of SVM-weighted neural ensembles

We analyzed the activity of push-weighted and pull-weighted neural ensembles that biased the SVM to predict push or pull.

- We averaged the activity over the neurons in the push-weighted and pull-weighted ensembles time-locked to force peak of 3g cross actions (Fig. 2K,L).
- We separately analyzed the average activity of the identified D1- and D2-SPNs within these ensembles (Fig. 2M,N).
- In Fig. 3K, we averaged ensemble activity in the 100ms before force peak of 3g cross actions.
- In Fig. 3L,M, we time-locked ensemble activity to holographic stimulation.

Effect of holographic stimulation on actions

We compared force in stimulation trials versus no-stimulation trials that immediately preceded stimulation trials. No-stimulation trials that occurred closer in time to the previous stimulation trial than the next stimulation trial were excluded. We analyzed force sampled at 1 kHz and low-pass filtered at 30 Hz (10th-order Butterworth filter), providing both high resolution visualization and de-noising.

Given that stimulation occurred with a random (but small) latency from 3g cross, we performed an alignment procedure to match the force of no-stimulation trials to stimulation trials in the time leading up to stimulation. We identified the time in the no-stimulation trial that best matched the following stimulation trial on 1) the force at time of stimulation ("target instantaneous force") and 2) the average force in the 100 ms window preceding time of stimulation ("target preceding force"). Concretely, we identified the time in each no-stimulation trial that minimized a cost, which was the sum of two terms: 1) the absolute difference between force at the time in the no-stimulation trial and the "target instantaneous force", and 2) the absolute difference between the average force in the 100 ms preceding the time in the no-stimulation trial and the "target preceding force." This alignment was performed on the raw 1 kHz force trace before filtering at 30 Hz.

We analyzed the difference between stimulation trials locked to stimulation time and no-stimulation trials locked to this aligned time. Trials were pooled into "congruent stimulation" and "non-congruent stimulation" conditions (Fig. 4). Congruent stimulation consisted of 1) stimulation of the push ensemble triggered on push 3g cross, and 2) stimulation of the pull ensemble triggered on pull 3g cross. Non-congruent stimulation consisted of 1) stimulation of the push ensemble triggered on pull 3g cross, and 2) stimulation of the pull ensemble triggered on push 3g cross. Accordingly, the number of data points for statistics (Fig. 4C,D) was 2×num_sessions.

Statistical analysis

Statistical analysis was performed using custom scripts in Python. We used the ‘scipy.stats’ package to perform paired t-tests (‘ttest_rel’ method), unpaired t-tests (‘ttest_ind’ method), and linear regression (‘linregress’ method). Linear mixed effects models were performed using the ‘statsmodels’ package and ‘mixedlm’ method. Significance was set at $P$=0.05. Statistical information in Tables S1 and S2.

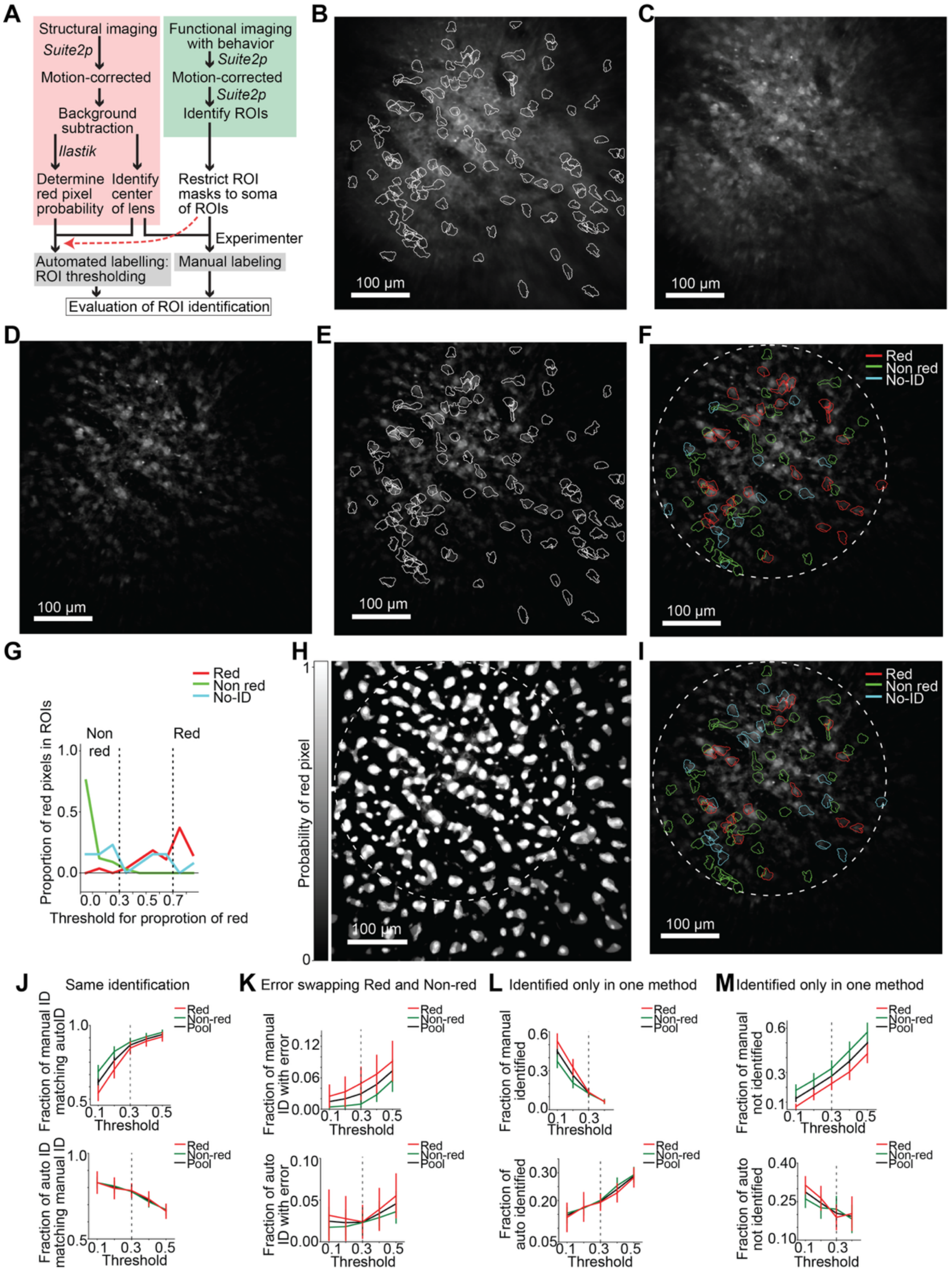

**Fig. S1. Method to label neuronal identity based on structural fluorescence.**
(**A**) Scheme of automated thresholding method to distinguish D1-SPNs from D2-SPNs. (**B**) Example of Suite2p identified ROIs (in white) overlaid on average image of functional channel (1000 frames of green channel). (**C**) Example of average image of structural channel (1000 frames of red channel). (**D**) Example of background subtracted structural image (rolling ball of 25 pixels). (**E**) Example of ROIs overlaid on structural image used for manual labeling. (**F**) Example of manual labeling ROIs within region of interest of GRIN lens (dashed white line). ROIs that experimenter couldn't confidently label are "No-ID." (**G**) For this example, distribution of red probability in manually labelled groups of ROIs (red for Red, green for no-

Red and cyan for No-ID). Dashed lines represent thresholds used for automated method results shown in (I). (**H**) Example of probability of pixel being red determined using machine learning software *Ilastik*. (**I**) Example of automated labeling ROIs within region of interest of GRIN lens (dashed white line). ROIs that automated method couldn't confidently label with thresholds are "No-ID." (**J-M**) Neuron identification accuracy as a function of automatic labeling threshold. ROIs with proportion of red pixels below threshold are labeled green, and ROIs with proportion of red pixels above $1 - $ threshold are labeled red. (**J**) Identification accuracy across thresholds. Top: fraction of ROIs identified manually that were also identified automatically. Bottom: fraction of ROIs identified automatically that were also identified manually. (**K**) Errors consisting of swapping "Red" and "Non-red" identity between identification methods. Top: the fraction of manually identified ROIs that were automatically identified with the opposite ID. Bottom: the fraction of automatically identified ROIs that were manually identified with the opposite ID. (**L**) Top: the fraction of manually identified ROIs that were not automatically identified. Bottom: the fraction of automatically identified ROIs that were not manually identified. (**M**) Top: the fraction of manually not-identified ROIs that were automatically identified. Bottom: the fraction automatically not-identified ROIs that were manually identified. Scale bar: 100 μm.

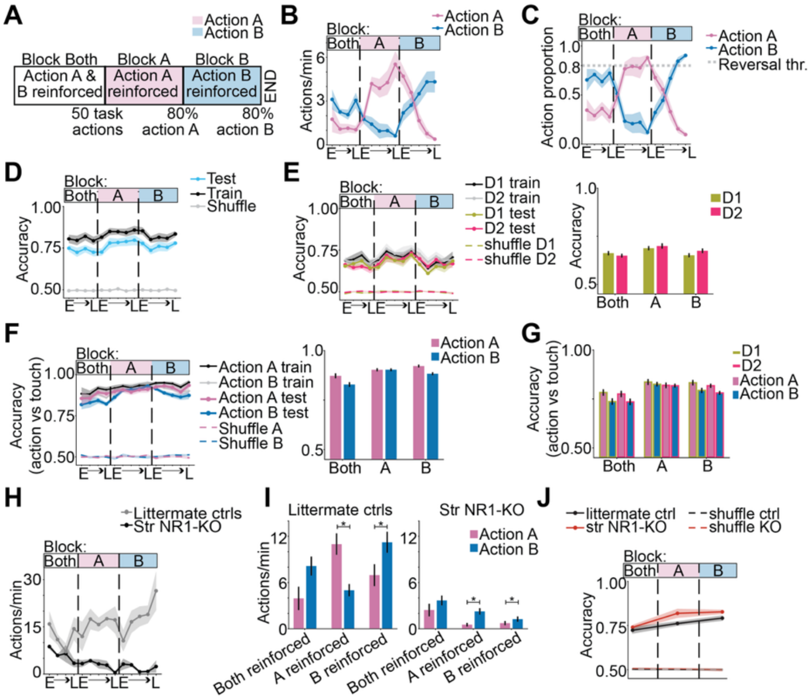


**Fig. S2. Striatal encoding of isometric actions irrespective of reinforcement or plasticity at striatal inputs.**

(**A**) Type or Schematic of reinforcement blocks. Criteria to advance to the next block are labelled. (**B**) The rate of task actions. In Block Both, the rate of Action B was greater than Action A (table S2, ref. 2.1). In Block A, Action A rate increased with sessions (table S2, ref. 2.2), and Action B rate decreased with sessions (table S2, ref. 2.3). In Block B, Action B rate increased with sessions (table S2, ref. 2.5), and Action A rate decreased with sessions (table S2, ref. 2.4). (**C**) The proportion of task actions from each identity (A or B). In Block A, the Action A proportion increased with sessions (table S2, ref. 2.6). In Block B, the Action B proportion increased with sessions (table S2, ref. 2.9). (**D**) Decoding SPN activity at force peak to predict action identity. Accuracy was greater than shuffle on the first session (table S2, ref. 2.10), in pooled sessions from Block Both, Block A and Block B (table S2, ref. 2.11, 2.12, 2.13). (**E**) Same as (D) but decoding D1- or D2-SPN activity. Right: average of sessions in each block. Accuracy was greater than shuffle in pooled sessions from Block Both, Block A and Block B (table S2; ref. for D1: 2.16, 2.17, 2.18; ref. for D2: 2.19, 2.20, 2.21). There was no significant difference between D1- and D2-SPNs in each block (table S2, ref. 2.22-2.24). (**F**) Decoding SPN activity at force peak to predict Action A from touch and Action B from touch. Right: average of sessions in each block. Accuracy was greater than shuffle on the first session (table S2, ref. 2.25, 2.26) and in pooled sessions from Block Both, Block A and Block B (table S2; ref. for Action A: 2.27, 2.28, 2.29; ref for Action B: 2.30, 2.31, 2.32). (**G**) Same as (F) Right but decoding D1- or D2-SPN activity. Decoding accuracy of Action A versus touch and Action B versus touch was greater than shuffle in pooled sessions from Block Both, Block A and Block B (table S2; ref. for D1: 2.37-2.42; ref for D2: 2.43-2.48). (**B-G**), Data are mean+s.e.m. across n=8 mice for each session or for each block in bar plots: 4 sessions for Block Both, 5 sessions for Block A and 5 sessions for Block B. (**H**) The rate of 6g cross actions (pooling push and pull) for control and mutant striatal NR1-KO mice. Data are mean+s.e.m. across n=4 control mice and n=3 NR1-KO mice for each session. For mutants, the action rate decreased with sessions (table S2, ref. 2.49).

For control mice, the action rate increased with sessions (table S2, ref. 2.50). (**I**) The rate of 6g cross actions in each block for control and mutant striatal NR1-KO mice. Littermate controls' rate for each action was not significantly different when both actions were reinforced (table S2, ref. 2.51). Littermate controls performed Action A more than Action B when Action A was reinforced (table S2, ref. 2.52) and Action B more than Action A when Action B was reinforced (table S2, ref. 2.53). NR1-KO mice' rate for each action was not significantly different when both actions were reinforced (table S2, ref. 2.54). NR1-KO mice performed Action B more than Action A when Action A was reinforced and when B was reinforced (table S2, ref. 2.55, 2.56, respectively). (**J**) Decoding SPN activity at force peak for littermate control and NR1-KO mice. Decoding accuracy was greater than shuffle in pooled sessions from Block Both, Block A and Block B (table S2, table ref. NR1-KO: 2.57, 2.58, 2.59; controls: 2.60, 2.61, 2.62). (**I-J**) Data are mean+s.e.m. over n=4 littermate control mice and n=3 NR1-KO mice, averaging sessions within each block.

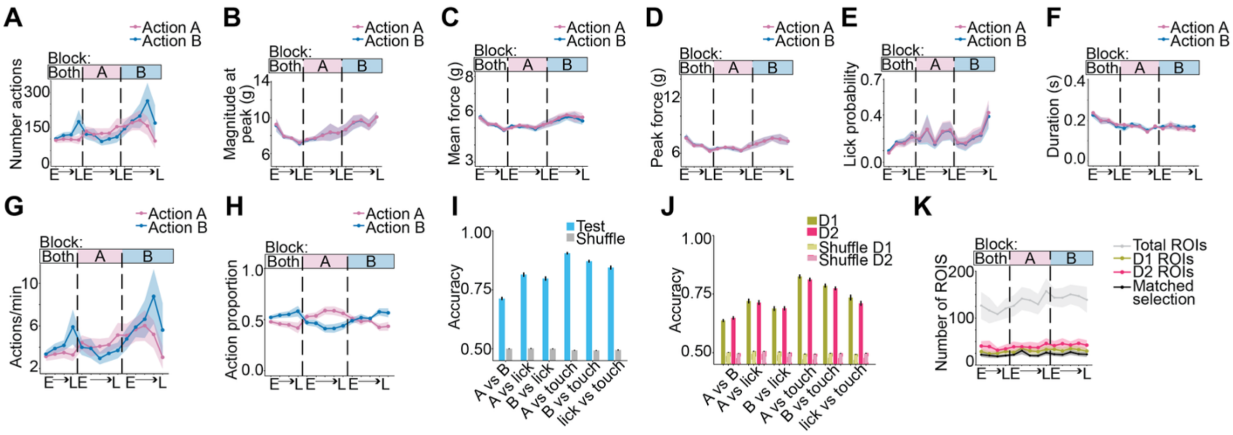

**Fig. S3. D1- and D2-SPNs encode the identity of isometric actions with matched continuous parameters.**

(**A**) Number of matched 3g cross actions per session. (**B**) Magnitude of two-dimensional force vector at time of peak force in push/pull axis. (**C**) Absolute force (in push/pull axis) averaged over action duration. (**D**) Peak force. (**E**) Lick probability averaged over duration. (**F**) Action duration. (**G**) Action rate. (**H**) Proportion of matched 3g cross actions with a specific identity (A or B). (**A-H**) Data are mean+s.e.m. for n=8 mice for each session. (**I**) Decoding SPN activity at force peak for matched 3g cross action A and B, predicting action identity between pairs of actions including A, B, touch, and licking. (**J**) Same as (I) but decoding D1- and D2-SPNs separately. (**K**) Number of functional ROIs identified with Suite2p. Decoders using D1- and D2-SPNs used ROIs with numbers matched across cell type (shown in black).

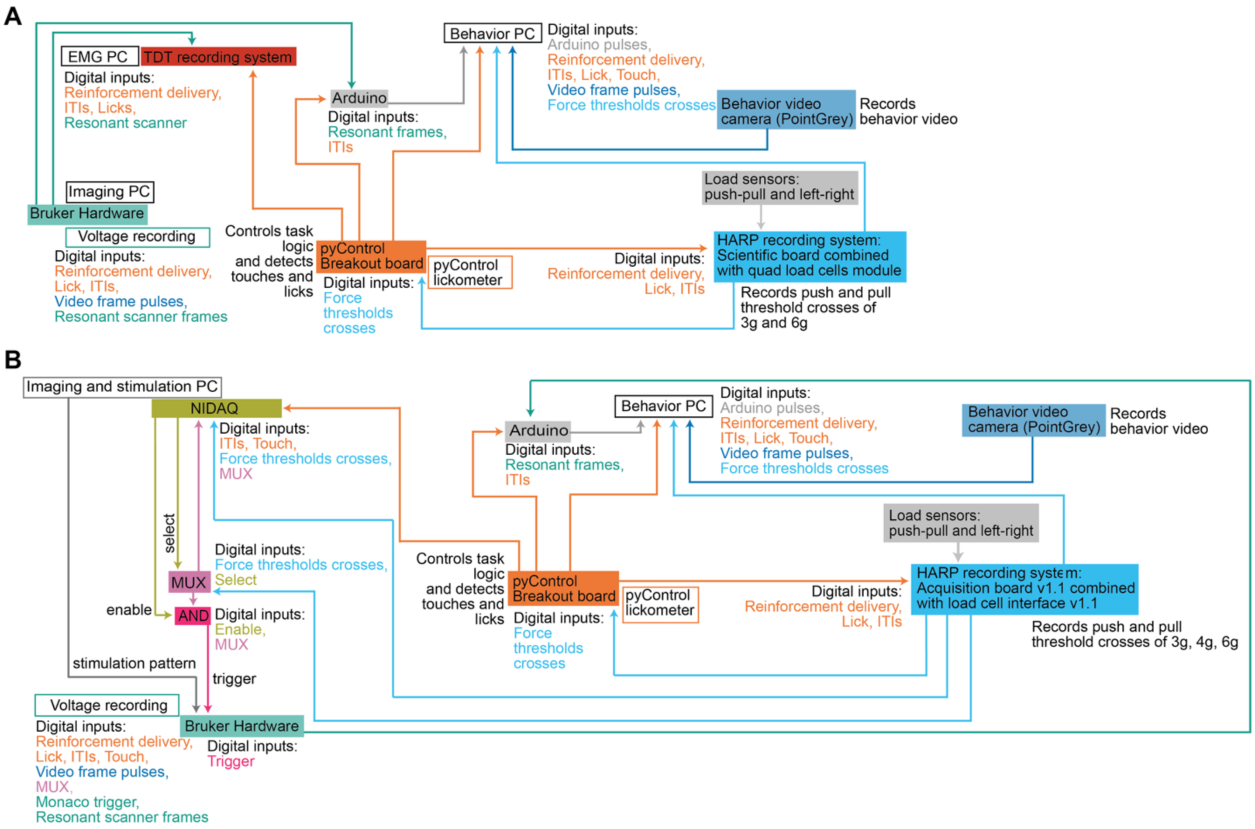


**Fig. S4. Experiment hardware setup schematics.**
(**A**) Schematic of hardware setup to run the two-action isometric task. (**B**) Schematic of hardware setup to run the adaptive isometric task and closed-loop holographic stimulation experiments.

| Table ref. | Figure Panel | Group | Statistical test | Sample size | Test Statistic | *P* value | Sig. | Notes |
|---|---|---|---|---|---|---|---|---|
| 1.1 | 1D | Action rate in Block Both: push vs pull | Paired t-test | 32 sessions (4 sessions/mouse, 8 mice) | 2.56 | 1.6e-2 | * | All sessions within block |
| 1.2 | 1D | Action rate in Block where Pull was reinforced: push vs pull | Paired t-test | 40 sessions (5 sessions/mouse, 8 mice) | 6.27 | 2.43e-7 | * | All sessions within block |
| 1.3 | 1D | Action rate in Block where Push was reinforced: push vs pull | Paired t-test | 40 sessions (5 sessions/mouse, 8 mice) | 3.38 | 1.65e-3 | * | All sessions within block |
| 1.4 | 1F | Action onset vs baseline: biceps | Paired t-test | 41 sessions (4 mice) | 5.47 | 2.61e-6 | * | 100 ms at 6g cross vs 100 ms at 1 s before 6g cross |
| 1.5 | 1F | Action onset vs baseline: triceps | Paired t-test | 41 sessions (4 mice) | 7.84 | 1.31e-9 | * | 100 ms at 6g cross vs 100 ms at 1 s before 6g cross |
| 1.6 | 1F | Action onset vs baseline: PL | Paired t-test | 41 sessions (4 mice) | 4.64 | 3.73e-5 | * | 100 ms at 6g cross vs 100 ms at 1 s before 6g cross |
| 1.7 | 1F | Action onset vs baseline: EDC | Paired t-test | 41 sessions (4 mice) | 3.68 | 6.8e-4 | * | 100 ms at 6g cross vs 100 ms at 1 s before 6g cross |
| 1.8 | 1G | Action onset vs baseline: biceps | Paired t-test | 41 sessions (4 mice) | 12.02 | 7.41e-15 | * | 100 ms at 6g cross vs 100 ms at 1 s before 6g cross |
| 1.9 | 1G | Action onset vs baseline: triceps | Paired t-test | 41 sessions (4 mice) | 6.76 | 4.11e-8 | * | 100 ms at 6g cross vs 100 ms at 1 s before 6g cross |
| 1.10 | 1G | Action onset vs baseline: PL | Paired t-test | 41 sessions (4 mice) | 7.39 | 5.37e-9 | * | 100 ms at 6g cross vs 100 ms at 1 s before 6g cross |
| 1.11 | 1G | Action onset vs baseline: EDC | Paired t-test | 41 sessions (4 mice) | 7.26 | 8.14e-9 | * | 100 ms at 6g cross vs 100 ms at 1 s before 6g cross |
| 1.12 | 1H | Difference across vs within action: push | Paired t-test | 41 sessions (4 mice) | 4.50 | 5.76e-5 | * | 100 ms centered at 6g cross |
| 1.13 | 1H | Difference across vs within action: pull | Paired t-test | 41 sessions (4 mice) | 6.60 | 6.87e-8 | * | 100 ms centered at 6g cross |
| 2.1 | 2C - push | Difference between baseline vs force peak: neural activity D1 | Paired t-test | 86 sessions (8 mice) | -9.19 | 2.06e-14 | * | ~90 ms (3 frames) at force peak and -0.5 s before peak |
| 2.2 | 2C - push | Difference between baseline vs force peak: neural activity D2 | Paired t-test | 86 sessions (8 mice) | 11.61 | 2.01e-14 | * | ~90 ms (3 frames) at force peak and -0.5 s before peak |
| 2.3 | 2C - pull | Difference between baseline vs force peak: neural activity D1 | Paired t-test | 86 sessions (8 mice) | -14.36 | 1.45e-24 | * | ~90 ms (3 frames) at force peak and -0.5 s before peak |

| | | | | | | | | |
|---|---|---|---|---|---|---|---|---|
| 2.4 | 2C - pull | Difference between baseline vs force peak: neural activity D2 | Paired t-test | 86 sessions (8 mice) | -14.78 | 2.51e-25 | * | ~90 ms (3 frames) at force peak and -0.5 s before peak |
| 2.5 | 2C - push | Difference between D1 and D2 at peak | Paired t-test | 86 sessions (8 mice) | 2.15 | 0.03 | * | -0.5 s to 0.5 s at force peak |
| 2.6 | 2C - pull | Difference between D1 and D2 at peak | Paired t-test | 86 sessions (8 mice) | 2.49 | 0.01 | * | -0.5 s to 0.5 s at force peak |
| 2.7 | 2E | Accuracy prediction at force peak: data vs shuffle | Paired t-test | 87 sessions (8 mice) | 25.85 | 2.55e-42 | * | 5 frames centered at peak force |
| 2.8 | 2E | Accuracy prediction before force peak: data vs shuffle | Paired t-test | 87 sessions (8 mice) | 11.45 | 5.50e-19 | * | Average over 4 sec before force peak |
| 2.9 | 2E | Accuracy prediction before force peak: data vs shuffle | Paired t-test | 87 sessions (8 mice) | 12.68 | 2.20e-21 | * | Average over 0.5 sec before force peak |
| 2.10 | 2G | Accuracy prediction at force peak: D1 data vs shuffle | Paired t-test | 87 sessions (8 mice) | 16.99 | 3.17e-29 | * | 5 frames centered at peak force |
| 2.11 | 2G | Accuracy prediction at force peak: D2 data vs shuffle | Paired t-test | 87 sessions (8 mice) | 16.24 | 6.18e-28 | * | 5 frames centered at peak force |
| 2.12 | 2G | Accuracy prediction at force peak: D2 data vs D1 data | Paired t-test | 87 sessions (8 mice) | -0.05 | 0.96 | n.s. | 5 frames centered at peak force |
| 2.13 | 2G | Accuracy prediction at force peak: D1 data vs shuffle | Paired t-test | 87 sessions (8 mice) | 8.91 | 7.49e-14 | * | 4 sec before force peak |
| 2.14 | 2G | Accuracy prediction at force peak: D2 data vs shuffle | Paired t-test | 87 sessions (8 mice) | 7.94 | 6.89e-12 | * | 4 sec before force peak |
| 2.15 | 2G | Accuracy prediction at force peak: D1 data vs shuffle | Paired t-test | 87 sessions (8 mice) | 10.33 | 9.72e-17 | * | 0.5 sec before force peak |
| 2.16 | 2G | Accuracy prediction at force peak: D2 data vs shuffle | Paired t-test | 87 sessions (8 mice) | 8.52 | 4.57e-13 | * | 0.5 sec before force peak |
| 2.17 | 2H | Accuracy prediction: push vs pull | Paired t-test | 86 sessions (8 mice) | 36.44 | 3.82e-62 | * | |
| 2.18 | 2H | Accuracy prediction: push vs lick | Paired t-test | 86 sessions (8 mice) | 26.29 | 1.21e-45 | * | |
| 2.19 | 2H | Accuracy prediction: pull vs lick | Paired t-test | 86 sessions (8 mice) | 28.99 | 2.81e-49 | * | |
| 2.20 | 2H | Accuracy prediction: push vs touch | Paired t-test | 86 sessions (8 mice) | 54.20 | 1.24e-79 | * | |
| 2.21 | 2H | Accuracy prediction: pull vs touch | Paired t-test | 86 sessions (8 mice) | 79.37 | 6.43e-97 | * | |
| 2.22 | 2H | Accuracy prediction: lick vs touch | Paired t-test | 86 sessions (8 mice) | 36.38 | 5.90e-58 | * | |

| | | | | | | | | |
|---|---|---|---|---|---|---|---|---|
| 2.23 | 2I | Accuracy predicting push vs pull: D1 vs D2 | Paired t-test | 86 sessions (8 mice) | -1.44 | 0.15 | n.s. | |
| 2.24 | 2I | D1 - Accuracy prediction: push vs pull | Paired t-test | 86 sessions (8 mice) | 24.38 | 6.49e-41 | * | |
| 2.25 | 2I | D1 - Accuracy prediction: push vs lick | Paired t-test | 86 sessions (8 mice) | 15.39 | 2.41e-25 | * | |
| 2.26 | 2I | D1 - Accuracy prediction: pull vs lick | Paired t-test | 86 sessions (8 mice) | 19.72 | 4.83e-32 | * | |
| 2.27 | 2I | D1 - Accuracy prediction: push vs touch | Paired t-test | 86 sessions (8 mice) | 28.51 | 3.16e-46 | * | |
| 2.28 | 2I | D1 - Accuracy prediction: pull vs touch | Paired t-test | 86 sessions (8 mice) | 38.42 | 6.89e-58 | * | |
| 2.29 | 2I | D1 - Accuracy prediction: lick vs touch | Paired t-test | 86 sessions (8 mice) | 19.14 | 3.35e-31 | * | |
| 2.30 | 2I | D2 - Accuracy prediction: push vs pull | Paired t-test | 86 sessions (8 mice) | 22.27 | 6.38e-38 | * | |
| 2.31 | 2I | D2 - Accuracy prediction: push vs lick | Paired t-test | 86 sessions (8 mice) | 16.28 | 8.32e-27 | * | |
| 2.32 | 2I | D2 - Accuracy prediction: pull vs lick | Paired t-test | 86 sessions (8 mice) | 18.47 | 3.24e-30 | * | |
| 2.33 | 2I | D2 - Accuracy prediction: push vs touch | Paired t-test | 86 sessions (8 mice) | 34.04 | 1.94e-52 | * | |
| 2.34 | 2I | D2 - Accuracy prediction: pull vs touch | Paired t-test | 86 sessions (8 mice) | 37.93 | 2.60e-56 | * | |
| 2.35 | 2I | D2 - Accuracy prediction: lick vs touch | Paired t-test | 86 sessions (8 mice) | 19.83 | 3.37e-32 | * | |
| 2.36 | 2J | Force peak SVM projection vs baseline: push | Paired t-test | 86 sessions (8 mice) | -10.75 | 1.36e-17 | * | -0.5 s to peak and at force peak |
| 2.37 | 2J | Force peak SVM projection vs baseline: pull | Paired t-test | 86 sessions (8 mice) | 17.05 | 2.57e-29 | * | -0.5 s to peak and at force peak |
| 2.38 | 2J | Force peak SVM projection pus vs pull | Paired t-test | 86 sessions (8 mice) | 25.33 | 1.21e-41 | * | -0.5 s to peak and at force peak |
| 2.39 | 2K | Push ensemble activity at peak vs before: push | Paired t-test | 86 sessions (8 mice) | -15.32 | 2.67e-26 | * | -0.5 s to peak and at force peak |
| 2.40 | 2K | Push ensemble activity at push peak vs pull peak | Paired t-test | 86 sessions (8 mice) | 19.33 | 4.70e-33 | * | At force peak |
| 2.41 | 2L | Pull ensemble activity at peak vs before: pull | Paired t-test | 86 sessions (8 mice) | -21.60 | 1.71e-36 | * | -0.5 s to peak and at force peak |
| 2.42 | 2L | Pull ensemble activity at push peak vs pull peak | Paired t-test | 86 sessions (8 mice) | -21.21 | 6.45e-36 | * | At force peak |
| 2.43 | 2M | Push ensemble activity at peak D1 vs D2 | Paired t-test | 63 sessions (8 mice) | 0.82 | 0.41 | n.s. | Average centered at force peak over -0.5s to 0.5s |
| 2.44 | 2N | Pull ensemble activity at peak D1 vs D2 | Paired t-test | 73 sessions (8 mice) | 0.68 | 0.50 | n.s. | Average centered at force peak over -0.5s to 0.5s |

| | | | | | | | | |
|---|---|---|---|---|---|---|---|---|
| 2.45 | Supplemental to Fig. 2 | Accuracy prediction: push versus pull at 3g cross; data vs shuffle | Paired t-test | 84 sessions (8 mice) | 13.08 | 3.28e-22 | * | Average of accuracy from -0.5 s to 0 s locked to 3g cross. Trials were analyzed with no other overlapping actions in this window. |
| 2.46 | Supplemental to Fig. 2 | Angle alignment (to force peak decoder) vs time (at which decoder is fit) | Linear regression | 86 sessions (8 mice) | -0.722 | 3.15e-113 | * | Testing if angle to force peak decoder decreases with time lags across the 0.5 seconds before 3g cross |
| 2.47 | Supplemental to Fig. 2 | Comparing angle alignment (to force peak decoder) at 3g cross versus at 0.467s before 3g cross | Paired t-test | 86 sessions (8 mice) | 34.50 | 3.79e-52 | * | |
| 2.48 | Supplemental to Fig. 2 | Comparing angle alignment (to force peak decoder) at 3g cross versus at 0.4s before 3g cross | Paired t-test | 86 sessions (8 mice) | 31.22 | 1.08e-48 | * | |
| 2.49 | Supplemental to Fig. 2 | Comparing angle alignment (to force peak decoder) at 3g cross versus at 0.33s before 3g cross | Paired t-test | 86 sessions (8 mice) | 30.30 | 1.16e-47 | * | |
| 2.50 | Supplemental to Fig. 2 | Comparing angle alignment (to force peak decoder) at 3g cross versus at 0.267s before 3g cross | Paired t-test | 86 sessions (8 mice) | 29.07 | 3.0e-46 | * | |
| 2.51 | Supplemental to Fig. 2 | Comparing angle alignment (to force peak decoder) at 3g cross versus at 0.2s before 3g cross | Paired t-test | 86 sessions (8 mice) | 29.61 | 7.06e-47 | * | |
| 2.52 | Supplemental to Fig. 2 | Comparing angle alignment (to force peak decoder) at 3g cross versus at 0.133s before 3g cross | Paired t-test | 86 sessions (8 mice) | 25.89 | 2.25e-42 | * | |
| 2.53 | Supplemental to Fig. 2 | Comparing angle alignment (to force peak decoder) at 3g cross versus at 0.067s before 3g cross | Paired t-test | 86 sessions (8 mice) | 25.81 | 2.91e-42 | * | |
| 3.1 | 3K | Push ensemble activity: push actions vs pull action | Paired t-test | 21 sessions (10 mice) | 5.58 | 1.86-5 | * | Average over 0.1 s before peak to peak in calibration |
| 3.2 | 3K | Pull ensemble activity: push actions vs pull action | Paired t-test | 21 sessions (10 mice) | -4.98 | 7.16e-5 | * | Average over 0.1 s before peak to peak in calibration |
| 3.3 | 3L | Neural activity: push ensemble vs pull ensemble | Paired t-test | 21 sessions (10 mice) | 5.35 | 3.06e-5 | * | Average over 0.1 s after stimulation |

| | | | | | | | | |
|---|---|---|---|---|---|---|---|---|
| 3.4 | 3L | Neural activity: push ensemble vs non-targeted | Paired t-test | 21 sessions (10 mice) | 5.55 | 1-97e-5 | * | Average over 0.1 s after stimulation |
| 3.5 | 3L | Neural activity: pull ensemble vs non-targeted | Paired t-test | 21 sessions (10 mice) | 0.24 | 0.81 | n.s. | Average over 0.1 s after stimulation |
| 3.6 | 3M | Neural activity: pull ensemble vs push ensemble | Paired t-test | 21 sessions (10 mice) | 4.33 | 3.26e-4 | * | Average over 0.1 s after stimulation |
| 3.7 | 3M | Neural activity: pull ensemble vs non-targeted | Paired t-test | 21 sessions (10 mice) | 4.48 | 2.30e-4 | * | Average over 0.1 s after stimulation |
| 3.8 | 3M | Neural activity: push ensemble vs non-targeted | Paired t-test | 21 sessions (10 mice) | 1.50 | 0.15 | n.s. | Average over 0.1 s after stimulation |
| 3.9 | 3N | SVM projection: post stim push ensemble | Paired t-test | 21 sessions (10 mice) | 3.53 | 2.10e-3 | * | Average over 0.1 s before stim. versus after stim. |
| 3.10 | 3N | SVM projection: post stim pull ensemble | Paired t-test | 21 sessions (10 mice) | 4.85 | 9.63e-5 | * | Average over 0.1 s before stim. versus after stim. |
| 3.11 | 3N | SVM projection: stim push ensemble vs pull ensemble | Paired t-test | 21 sessions (10 mice) | 4.90 | 8.57e-5 | * | Average over 0.1 s before stim. versus after stim. |
| 4.1 | 4C | D1 – force perturbation: congruent vs non-congruent: 0 to 0.01 s | One-sided paired t-test | 24 session-conditions (12*2) | 5.75e-1 | 2.85e-1 | n.s. | Selected interval locked to stimulation onset |
| 4.2 | 4C | D1 – force perturbation: congruent vs non-congruent: 0.01 to 0.02 s | One-sided paired t-test | 24 session-conditions (12*2) | 6.80e-1 | 2.52e-1 | n.s. | Selected interval locked to stimulation onset |
| 4.3 | 4C | D1 – force perturbation: congruent vs non-congruent: 0.02 to 0.03 s | One-sided paired t-test | 24 session-conditions (12*2) | 1.10 | 1.40-1 | n.s. | Selected interval locked to stimulation onset |
| 4.4 | 4C | D1 – force perturbation: congruent vs non-congruent: 0.03 to 0.04 s | One-sided paired t-test | 24 session-conditions (12*2) | 1.64 | 5.77e-2 | n.s. | Selected interval locked to stimulation onset |
| 4.5 | 4C | D1 – force perturbation: congruent vs non-congruent: 0.04 to 0.05 s | One-sided paired t-test | 24 session-conditions (12*2) | 2.15 | 2.10e-2 | * | Selected interval locked to stimulation onset |
| 4.6 | 4C | D1 – force perturbation: congruent vs non-congruent: 0.05 to 0.06 s | One-sided paired t-test | 24 session-conditions (12*2) | 2.45 | 1.11e-2 | * | Selected interval locked to stimulation onset |
| 4.7 | 4C | D1 – force perturbation: congruent vs non-congruent: 0.06 to 0.07 s | One-sided paired t-test | 24 session-conditions (12*2) | 2.54 | 9.07e-3 | * | Selected interval locked to stimulation onset |
| 4.8 | 4C | D1 – force perturbation: congruent vs non-congruent: 0.07 to 0.08 s | One-sided paired t-test | 24 session-conditions (12*2) | 2.65 | 7.23e-3 | * | Selected interval locked to stimulation onset |
| 4.9 | 4C | D1 – force perturbation: congruent vs non-congruent: 0.08 to 0.09 s | One-sided paired t-test | 24 session-conditions (12*2) | 2.78 | 5.28e-3 | * | Selected interval locked to stimulation onset |

| | | | | | | | | |
|---|---|---|---|---|---|---|---|---|
| 4.10 | 4C | D1 – force perturbation: congruent vs non-congruent: 0.09 to 0.10 s | One-sided paired t-test | 24 session-conditions (12*2) | 2.78 | 5.31e-3 | * | Selected interval locked to stimulation onset |
| 4.11 | 4C | D1 – force perturbation: congruent vs non-congruent: 0.10 to 0.11 s | One-sided paired t-test | 24 session-conditions (12*2) | 2.51 | 9.85e-3 | * | Selected interval locked to stimulation onset |
| 4.12 | 4C | D1 – force perturbation: congruent vs no stimulation: 0.10 to 0.11 s | One-sided t-test | 24 session-conditions (12*2) | 2.29 | 1.57e-2 | * | Selected interval locked to stimulation onset |
| 4.13 | 4C | D1 – force perturbation: non-congruent vs no stimulation: 0.10 to 0.11 s | One-sided t-test | 24 session-conditions (12*2) | -0.30 | 6.15e-1 | n.s. | Selected interval locked to stimulation onset |
| 4.14 | 4D | D2 – force perturbation: congruent vs non-congruent: 0 to 0.01 s | One-sided paired t-test | 18 session-conditions (9*2) | 7.22e-1 | 2.40e-1 | n.s. | Selected interval locked to stimulation onset |
| 4.15 | 4D | D2 – force perturbation: congruent vs non-congruent: 0.01 to 0.02 s | One-sided paired t-test | 18 session-conditions (9*2) | 2.27.-1 | 4.11e-1 | n.s. | Selected interval locked to stimulation onset |
| 4.16 | 4D | D2 – force perturbation: congruent vs non-congruent: 0.02 to 0.03 s | One-sided paired t-test | 18 session-conditions (9*2) | 1.76e-1 | 4.31e-1 | n.s. | Selected interval locked to stimulation onset |
| 4.17 | 4D | D2 – force perturbation: congruent vs non-congruent: 0.03 to 0.04 s | One-sided paired t-test | 18 session-conditions (9*2) | 3.68e-1 | 3.59e-1 | n.s. | Selected interval locked to stimulation onset |
| 4.18 | 4D | D2 – force perturbation: congruent vs non-congruent: 0.04 to 0.05 s | One-sided paired t-test | 18 session-conditions (9*2) | 6.45e-1 | 2.65e-1 | n.s. | Selected interval locked to stimulation onset |
| 4.19 | 4D | D2 – force perturbation: congruent vs non-congruent: 0.05 to 0.06 s | One-sided paired t-test | 18 session-conditions (9*2) | 7.96e-1 | 2.19e-1 | n.s. | Selected interval locked to stimulation onset |
| 4.20 | 4D | D2 – force perturbation: congruent vs non-congruent: 0.06 to 0.07 s | One-sided paired t-test | 18 session-conditions (9*2) | 8.41e-1 | 2.06e-1 | n.s. | Selected interval locked to stimulation onset |
| 4.21 | 4D | D2 – force perturbation: congruent vs non-congruent: 0.07 to 0.08 s | One-sided paired t-test | 18 session-conditions (9*2) | 9.48e-1 | 1.78e-1 | n.s. | Selected interval locked to stimulation onset |
| 4.22 | 4D | D2 – force perturbation: congruent vs non-congruent: 0.08 to 0.09 s | One-sided paired t-test | 18 session-conditions (9*2) | 1.18 | 1.28e-1 | n.s. | Selected interval locked to stimulation onset |
| 4.23 | 4D | D2 – force perturbation: congruent vs non-congruent: 0.09 to 0.10 s | One-sided paired t-test | 18 session-conditions (9*2) | 1.49 | 7.78e-2 | n.s. | Selected interval locked to stimulation onset |
| 4.24 | 4D | D2 – force perturbation: congruent vs | One-sided paired t-test | 18 session-conditions (9*2) | 1.78 | 4.70e-2 | * | Selected interval locked to stimulation onset |

| | | | | | | | | |
|---|---|---|---|---|---|---|---|---|
| | | non-congruent: 0.10 to 0.11 s | | | | | | |
| 4.25 | 4D | D2 – force perturbation: congruent vs no stimulation: 0.10 to 0.11 s | One-sided t-test | 18 session-conditions (9*2) | 1.16 | 3.03e-2 | * | Selected interval locked to stimulation onset |
| 4.26 | 4D | D2 – force perturbation: non-congruent vs no stimulation: 0.10 to 0.11 s | One-sided t-test | 18 session-conditions (9*2) | 0.33 | 3.74e-1 | n.s. | Selected interval locked to stimulation onset |
| 4.27 | 4C | D1 – force perturbation; Congruent vs 0: 0 to 0.05 s | One-sided t-test | 24 session-conditions (12*2) | 2.24 | 1.74e-2 | * | Selected interval locked to stimulation onset |
| 4.28 | 4C | D1 – force perturbation; Congruent vs 0: 0.05 to 0.1 s | One-sided t-test | 24 session-conditions (12*2) | 2.58 | 8.36e-3 | * | Selected interval locked to stimulation onset |
| 4.29 | 4C | D1 – force perturbation; Congruent vs 0: 0.1 to 0.15 s | One-sided t-test | 24 session-conditions (12*2) | 1.96 | 3.08e-2 | * | Selected interval locked to stimulation onset |
| 4.30 | 4C | D1 – force perturbation; Non-congruent vs 0: 0 to 0.05 s | One-sided t-test | 24 session-conditions (12*2) | 1.09 | 1.44e-1 | n.s. | Selected interval locked to stimulation onset |
| 4.31 | 4C | D1 – force perturbation; Non-congruent vs 0: 0.05 to 0.1 s | One-sided t-test | 24 session-conditions (12*2) | -6.19e-2 | 5.24e-1 | n.s. | Selected interval locked to stimulation onset |
| 4.32 | 4C | D1 – force perturbation; Non-congruent vs 0: 0.1 to 0.15 s | One-sided t-test | 24 session-conditions (12*2) | 3.57e-2 | 4.86e-1 | n.s. | Selected interval locked to stimulation onset |
| 4.33 | 4D | D2 – force perturbation; Congruent vs 0: 0 to 0.05 s | One-sided t-test | 18 session-conditions (9*2) | 2.63 | 8.76e-3 | * | Selected interval locked to stimulation onset |
| 4.34 | 4D | D2 – force perturbation; Congruent vs 0: 0.05 to 0.1 s | One-sided t-test | 18 session-conditions (9*2) | 1.87 | 3.92e-2 | * | Selected interval locked to stimulation onset |
| 4.35 | 4D | D2 – force perturbation; Congruent vs 0: 0.1 to 0.15 s | One-sided t-test | 18 session-conditions (9*2) | 2.07 | 2.71e-2 | * | Selected interval locked to stimulation onset |
| 4.36 | 4D | D2 – force perturbation; Non-congruent vs 0: 0 to 0.05 s | One-sided t-test | 18 session-conditions (9*2) | 2.29 | 1.74e-2 | * | Selected interval locked to stimulation onset |
| 4.37 | 4D | D2 – force perturbation; Non-congruent vs 0: 0.05 to 0.1 s | One-sided t-test | 18 session-conditions (9*2) | 0.967 | 1.74e-1 | n.s. | Selected interval locked to stimulation onset |
| 4.38 | 4D | D2 – force perturbation; Non-congruent vs 0: 0.1 to 0.15 s | One-sided t-test | 18 session-conditions (9*2) | -0.173 | 5.67e-1 | n.s. | Selected interval locked to stimulation onset |

**Table S1. Summary of statistical tests in Figs. 1-4.**

| Table ref. | Figure Panel | Group | Statistical test | Sample size | Test Statistic | *P* value | Sig. | Notes |
|---|---|---|---|---|---|---|---|---|
| 2.1 | S2B | Action rate at Both reinforced: A vs B | Paired t-test | 32 sessions (8 mice) | -3.36 | 2.11e-3 | * | |
| 2.2 | S2B | Modeled Rate of action as dependent on session: Block A action A | Linear mixed effect model | 40 sessions (8 mice) | Slope: 0.74 | 2.6e-5 | * | Data grouped by animal; modeled as random effect |
| 2.3 | S2B | Modeled Rate of action as dependent on session: Block A action B | Linear mixed effect model | 40 sessions (8 mice) | Slope: -0.29 | 2.1e-3 | * | Data grouped by animal; modeled as random effect Negative slope indicates drops with sessions |
| 2.4 | S2B | Modeled Rate of action as dependent on session: Block B action A | Linear mixed effect model | 40 sessions (8 mice) | Slope: -1.14 | 1.2e-14 | * | Data grouped by animal; modeled as random effect Negative slope indicates drops with sessions |
| 2.5 | S2B | Modeled Rate of action as dependent on session: Block B action B | Linear mixed effect model | 40 sessions (8 mice) | Slope: 0.59 | 1.2e-14 | * | Data grouped by animal; modeled as random effect |
| 2.6 | S2C | Modeled Proportion of action as dependent on session: Block A action A | Linear mixed effect model | 40 sessions (8 mice) | Slope: 0.09 | 4.11e-4 | * | Data grouped by animal; modeled as random effect |
| 2.7 | S2C | Modeled Proportion of action as dependent on session: Block A action B | Linear mixed effect model | 40 sessions (8 mice) | Slope: -0.09 | 4.1e-4 | * | Data grouped by animal; modeled as random effect Negative slope indicates drops with sessions |
| 2.8 | S2C | Modeled Proportion of action as dependent on session: Block B action A | Linear mixed effect model | 40 sessions (8 mice) | Slope: -0.16 | 2.0e-17 | * | Data grouped by animal; modeled as random effect Negative slope indicates drops with sessions |
| 2.9 | S2C | Modeled Proportion of action as dependent on session: Block B action B | Linear mixed effect model | 40 sessions (8 mice) | Slope: 0.16 | 2.0e-17 | * | Data grouped by animal; modeled as random effect |
| 2.10 | S2D | 1st session: Accuracy predicting action identity: test vs shuffle | Paired t-test | 8 sessions (8 mice) | 7.40 | 1.5e-4 | * | |
| 2.11 | S2D | Block Both: Accuracy predicting action identity: test vs shuffle | Paired t-test | 32 sessions (8 mice) | 18.15 | 9.95e-18 | * | All sessions in block |
| 2.12 | S2D | Block A: Accuracy predicting action identity: test vs shuffle | Paired t-test | 40 sessions (8 mice) | 21.50 | 3.92e-22 | * | All sessions in block |

| | | | | | | | | |
|---|---|---|---|---|---|---|---|---|
| 2.13 | S2D | Block B: Accuracy predicting action identity: test vs shuffle | Paired t-test | 40 sessions (8 mice) | 23.90 | 6.73e-25 | * | All sessions in block |
| 2.14 | S2D | Accuracy predicting action identity: block Both vs Block A | Independent t-test | 72 sessions (8 mice) | 2.18 | 3.29e-2 | * | All sessions in block |
| 2.15 | S2D | Accuracy predicting action identity: block Both vs Block B | Independent t-test | 72 sessions (8 mice) | 1.64 | 1.05e-1 | n.s. | All sessions in block |
| 2.16 | S2E | D1- Block Both: Accuracy predicting action identity: test vs shuffle | Paired t-test | 32 sessions (8 mice) | 13.83 | 3.36e-13 | * | All sessions in block |
| 2.17 | S2E | D1 - Block A: Accuracy predicting action identity: test vs shuffle | Paired t-test | 40 sessions (8 mice) | 15.64 | 5.76e-16 | * | All sessions in block |
| 2.18 | S2E | D1 - Block B: Accuracy predicting action identity: test vs shuffle | Paired t-test | 40 sessions (8 mice) | 13.28 | 2.48e-14 | * | All sessions in block |
| 2.19 | S2E | D2- Block Both: Accuracy predicting action identity: test vs shuffle | Paired t-test | 32 sessions (8 mice) | 13.39 | 6.62e-13 | * | All sessions in block |
| 2.20 | S2E | D2 - Block A: Accuracy predicting action identity: test vs shuffle | Paired t-test | 40 sessions (8 mice) | 13.42 | 3.19e-14 | * | All sessions in block |
| 2.21 | S2E | D2 - Block B: Accuracy predicting action identity: test vs shuffle | Paired t-test | 40 sessions (8 mice) | 13.65 | 1.18e-14 | * | All sessions in block |
| 2.22 | S2E; bar plot | Block Both: D1 vs D2 accuracy difference | Paired t-test | 32 sessions (8 mice) | 1.46 | 1.57e-1 | n.s. | All sessions in block |
| 2.23 | S2E; bar plot | Block A: D1 vs D2 accuracy difference | Paired t-test | 40 sessions (8 mice) | -1.34 | 1.89e-1 | n.s. | All sessions in block |
| 2.24 | S2E; bar plot | Block B: D1 vs D2 accuracy difference | Paired t-test | 40 sessions (8 mice) | -2.01 | 5.28e-2 | n.s. | All sessions in block |
| 2.25 | S2F | Action A - 1st session: Accuracy predicting action identity | Paired t-test | 8 sessions (8 mice) | 15.35 | 1.20e-6 | * | |

| | | | | | | | | |
|---|---|---|---|---|---|---|---|---|
| | | vs touch: test vs shuffle | | | | | | |
| 2.26 | S2F | Action B - 1st session: Accuracy predicting action identity vs touch: test vs shuffle | Paired t-test | 8 sessions (8 mice) | 10.28 | 1.79e-5 | * | |
| 2.27 | S2F | Action A - Block Both: Accuracy predicting action identity vs touch: test vs shuffle | Paired t-test | 32 sessions (8 mice) | 28.27 | 3.51e-23 | * | All sessions in block |
| 2.28 | S2F | Action A - Block A: Accuracy predicting action identity vs touch: test vs shuffle | Paired t-test | 40 sessions (8 mice) | 44.68 | 3.92e-33 | * | All sessions in block |
| 2.29 | S2F | Action A - Block B: Accuracy predicting action identity vs touch: test vs shuffle | Paired t-test | 40 sessions (8 mice) | 50.05 | 5.32e-37 | * | All sessions in block |
| 2.30 | S2F | Action B - Block Both: Accuracy predicting action identity vs touch: test vs shuffle | Paired t-test | 32 sessions (8 mice) | 28.37 | 3.15e-23 | * | All sessions in block |
| 2.31 | S2F | Action B - Block A: Accuracy predicting action identity vs touch: test vs shuffle | Paired t-test | 40 sessions (8 mice) | 49.75 | 8.71e-35 | * | All sessions in block |
| 2.32 | S2F | Action B - Block B: Accuracy predicting action identity vs touch: test vs shuffle | Paired t-test | 40 sessions (8 mice) | 41.81 | 5.21e-34 | * | All sessions in block |
| 2.33 | S2F | Action A vs Touch - Accuracy predicting action identity vs touch: Block Both vs Block A | Independent t-test | 72 sessions (8 mice) | 2.04 | 4.58e-2 | * | All sessions in block |
| 2.34 | S2F | Action B vs Touch - Accuracy predicting action identity vs touch: Block Both vs Block A | Independent t-test | 72 sessions (8 mice) | 5.58 | 4.79e-7 | * | All sessions in block |
| 2.35 | S2F | Action A vs Touch - Accuracy predicting action identity vs touch: | Independent t-test | 72 sessions (8 mice) | 3.56 | 6.81e-4 | * | All sessions in block |

| | | | | | | | | |
|---|---|---|---|---|---|---|---|---|
| | | Block Both vs Block B | | | | | | |
| 2.36 | S2F | Action B vs Touch - Accuracy predicting action identity vs touch: Block Both vs Block B | Independent t-test | 72 sessions (8 mice) | 3.98 | 1.70e-4 | * | All sessions in block |
| 2.37 | S2G | D1: Action A - Block Both: Accuracy predicting action identity vs touch: test vs shuffle | Paired t-test | 32 sessions (8 mice) | 16.13 | 1.01e-14 | * | All sessions in block |
| 2.38 | S2G | D1: Action A - Block A: Accuracy predicting action identity vs touch: test vs shuffle | Paired t-test | 40 sessions (8 mice) | 21.96 | 4.84e-20 | * | All sessions in block |
| 2.39 | S2G | D1: Action A - Block B: Accuracy predicting action identity vs touch: test vs shuffle | Paired t-test | 40 sessions (8 mice) | 21.97 | 1.84e-20 | * | All sessions in block |
| 2.40 | S2G | D1: Action B - Block Both: Accuracy predicting action identity vs touch: test vs shuffle | Paired t-test | 32 sessions (8 mice) | 13.22 | 8.73e-13 | * | All sessions in block |
| 2.41 | S2G | D1: Action B - Block A: Accuracy predicting action identity vs touch: test vs shuffle | Paired t-test | 40 sessions (8 mice) | 24.79 | 1.54e-21 | * | All sessions in block |
| 2.42 | S2G | D1: Action B - Block B: Accuracy predicting action identity vs touch: test vs shuffle | Paired t-test | 40 sessions (8 mice) | 21.73 | 2.54e-20 | * | All sessions in block |
| 2.43 | S2G | D2: Action A - Block Both: Accuracy predicting action identity vs touch: test vs shuffle | Paired t-test | 32 sessions (8 mice) | 15.95 | 1.30e-14 | * | All sessions in block |
| 2.44 | S2G | D2: Action A - Block A: Accuracy predicting action identity vs touch: test vs shuffle | Paired t-test | 40 sessions (8 mice) | 22.54 | 2.33e-20 | * | All sessions in block |
| 2.45 | S2G | D2: Action A - Block B: Accuracy predicting action identity vs touch: test vs shuffle | Paired t-test | 40 sessions (8 mice) | 29.44 | 3.18e-24 | * | All sessions in block |

| | | | | | | | | |
|---|---|---|---|---|---|---|---|---|
| 2.46 | S2G | D2: Action B - Block Both: Accuracy predicting action identity vs touch: test vs shuffle | Paired t-test | 32 sessions (8 mice) | 14.05 | 2.29e-13 | * | All sessions in block |
| 2.47 | S2G | D2: Action B - Block A: Accuracy predicting action identity vs touch: test vs shuffle | Paired t-test | 40 sessions (8 mice) | 28.35 | 3.24e-23 | * | All sessions in block |
| 2.48 | S2G | D2: Action B - Block B: Accuracy predicting action identity vs touch: test vs shuffle | Paired t-test | 40 sessions (8 mice) | 25.05 | 3.92e-22 | * | All sessions in block |
| 2.49 | S2H, NR1-KO | Modeled total action rate as dependent on session | Linear mixed effect model | 12 sessions (3 mice) | Slope: -0.44 | 3.85e-8 | * | Data grouped by animal; modeled as random effect Negative slope indicates drop with sessions |
| 2.50 | S2H, littermate controls | Modeled total action rate as dependent on session | Linear mixed effect model | 20 sessions (4 mice) | Slope: 0.74 | 4.35e-3 | * | Data grouped by animal; modeled as random effect |
| 2.51 | S2I, littermate controls | Action rate at Both reinforced block: A vs B | Paired t-test | 20 sessions (4 mice) | -1.92 | 0.074 | n.s. | All sessions in block |
| 2.52 | S2I, littermate controls | Action rate at A reinforced block: A vs B | Paired t-test | 25 sessions (4 mice) | 3.72 | 1.45e-3 | * | All sessions in block |
| 2.53 | S2I, littermate controls | Action rate at B reinforced block: A vs B | Paired t-test | 25 sessions (4 mice) | -2.29 | 0.033 | * | All sessions in block |
| 2.54 | S2I, NR1-KO | Action rate at Both reinforced block: A vs B | Paired t-test | 12 sessions (3 mice) | -1.00 | 0.34 | n.s. | All sessions in block |
| 2.55 | S2I, NR1-KO | Action rate at A reinforced block: A vs B | Paired t-test | 15 sessions (3 mice) | -4.79 | 2.89e-4 | * | All sessions in block More B than A |
| 2.56 | S2I, NR1-KO | Action rate at B reinforced block: A vs B | Paired t-test | 15 sessions (3 mice) | -2.70 | 0.017 | * | All sessions in block More B than A |
| 2.57 | S2J, NR1-KO | NR1-KO - Block Both: Accuracy predicting action identity: test vs shuffle | Paired t-test | 12 sessions (3 mice) | 19.16 | 2.63e-7 | * | All sessions in block |
| 2.58 | S2J, NR1-KO | NR1-KO - Block A: Accuracy predicting action identity: test vs shuffle | Paired t-test | 15 sessions (3 mice) | 11.24 | 9.72e-5 | * | All sessions in block |
| 2.59 | S2J, NR1-KO | NR1-KO - Block B: Accuracy predicting action | Paired t-test | 15 sessions (3 mice) | 18.03 | 2.26e-8 | * | All sessions in block |

| | | | | | | | | |
|---|---|---|---|---|---|---|---|---|
| | | identity: test vs shuffle | | | | | | |
| 2.60 | S2J, littermate controls | Controls - Block Both: Accuracy predicting action identity: test vs shuffle | Paired t-test | 20 sessions (4 mice) | 14.27 | 9.80e-10 | * | All sessions in block |
| 2.61 | S2J, littermate controls | Controls - Block A: Accuracy predicting action identity: test vs shuffle | Paired t-test | 25 sessions (4 mice) | 25.36 | 1.54e-15 | * | All sessions in block |
| 2.62 | S2J, littermate controls | Controls - Block B: Accuracy predicting action identity: test vs shuffle | Paired t-test | 25 sessions (4 mice) | 24.38 | 8.49e-16 | * | All sessions in block |
| 2.63 | S2J | Accuracy predicting action identity: NR1-KO vs controls | Independent t-test | 112 sessions (4+3 animals) | 2.29 | 2.48e-2 | * | All sessions |

### Table S2. Summary of statistical tests in Fig. S2

| Line name | Extended name | Source | Available at | Code | Ref. |
|---|---|---|---|---|---|
| Tg(Drd1-cre)EY217Gsat | STOCK Tg(Drd1-cre)EY217Gsat/Mmucd | Gerfen lab* | MMRRC | 030778 | Gong et al, 2007(*53*) |
| Ai9(RCL-tdT) | B6;129S6-Gt(ROSA)26Sortm9(CAG-tdTomato)Hze/J | Jackson Laboratories | Jackson Laboratories | 007905 | Madisen et al, 2010(*52*) |
| Drd1a-tdTomato line 6 | B6.Cg-Tg(Drd1a-tdTomato)6Calak/J | Jackson Laboratories | Jackson Laboratories | 016204 | Ade et al, 2011(*51*) |
| Tg(Adora2a-cre)KG139Gsat | B6.FVB(Cg)-Tg(Adora2a-cre)KG139Gsat/Mmucd | MMRRC* | MMRRC | 036158 | Gong et al, 2007(*53*) |
| RGS9-cre | B6;129S-Rgs9$^{tm1.1(cre)Yql}$/J | NIH from Yuqing Li* | Jackson Laboratories | 020550 | Dang et al, 2006(*56*) |
| NMDAR1-loxP | B6.129S2(Cg)-*Grin1*$^{tm1Yql}$/NkzaJ | NIH from Yuqing Li* | Jackson Laboratories | 036352 | Dang et al, 2006(*56*) |

* These lines were kept in house and backcrossed for more than 10 generations.

**Table S3. Mouse lines used in experiments.**

| Reinforcement block | Parameters label | Task action threshold (g) | Task action duration (ms) | Reinforcement delay (ms) |
|---|---|---|---|---|
| Both | 1 | 4 | 10 | 100 |
| Both | 2 | 4 | 20 | 200 |
| Both | 3 | 6 | 20 | 300 |
| A or B | 1 | 4 | 20 | 300 |
| A or B | 2 | 6 | 20 | 300 |
| A or B | 3 | 6 | 50 | 300 |

**Table S4. Changes in task parameters for different reinforcement blocks.**

| Reinforcement block | Selected session | Selection Rules |
|---|---|---|
| Both | 1 | 1st day at Parameters 1 for Block Both |
| Both | 2 | 1st day at Parameters 2 for Block Both |
| Both | 3 | 1st day at Parameters 3 from Block Both |
| Both | 4 | Last day before change in reinforcement schedule |
| A or B | 1 | 1st day at Parameters 1 for Block A or B |
| A or B | 2 | 1st day at Parameters 2 or Block A or B |
| A or B | 3 | 1st day at Parameters 3 or Block A or B |
| A or B | 4 | Middle day between selected session 3 and 5 |
| A or B | 5 | Last day of current reinforcement Block |

**Table S5. Selected sessions for comparison across animals for all reinforcement blocks.**

| REAGENT or RESOURCE | SOURCE | IDENTIFIER |
|---|---|---|
| **Antibodies** | | |
| anti-GFP Alexa 488 conjugate | Molecular Probes | #A-21311 ; RRID:AB_221477 |
| rabbit anti-RFP | Rockland | 600-401-379; RRID:AB_828391 |
| anti-Rabbit Alexa Fluor 568 | Invitrogen | A11011; RRID:AB_143157 |
| **Bacterial and virus strains** | | |
| AAV5.CaMKII.GCaMP6f.WPRE.SV40 | Addgene | 100834-AAV5; RRID:Addgene_100834; lots: #vv59618 and t# v102673 |
| AAV8.CaMKII.ChRmine.mScarlet.Kv2.1.WPRE | Deisseroth lab; Marshel et al, 2019(*30*) | KD5172 |
| **Deposited data** | | |
| Histological images | This paper | https://doi.org/10.35077/g.1203 |
| 2-photon imaging | This paper | 10.6019/S-BIAD3134 |
| Behavior dataset | This paper | 10.6019/S-BIAD3134 |
| 2-photon imaging and stimulation | This paper | 10.6019/S-BIAD3135 |
| **Experimental models: Mouse strains** | | |
| Tg(Drd1-cre)EY217Gsat | MMRRC | 30778; RRID:MGI:4366805 |
| Ai9(RCL-tdT) | The Jackson Laboratory | 7905; RRID:IMSR_JAX:007905 |
| Drd1a-tdTomato line 6 | The Jackson Laboratory | 74016204; RRID:IMSR_JAX:016204 |
| Tg(Adora2a-cre)KG139Gsat | MMRRC | 36158; RRID:MMRRC_036158-UCD |
| RGS9-cre | The Jackson Laboratory | 20550; RRID:IMSR_JAX:020550 |
| NMDAR1-loxP | The Jackson Laboratory | 36352; RRID:IMSR_JAX:036352 |
| **Software and algorithms** | | |
| Python | | RRID:SCR_008394 |
| MATLAB | | RRID:SCR_001622 |
| pyControl | Akam et al, 2022(*75*) | RRID:SCR_021612 |
| statsmodels | | RRID:SCR_016074 |
| Suite2p | Pachitariu et al, 2017(*76*) | RRID:SCR_016434 |
| Prairie View | Bruker | RRID:SCR_017142 |
| Synapse | Tucker-Davis Technologies | RRID:SCR_006495 |
| Ilastik | Sommer et al, 2011(*78*) | RRID:SCR_015246 |
| ImageJ | | RRID:SCR_003070 |
| BrainJ | Botta et al, 2020(*73*) | RRID:SCR_027061 |
| Data extraction | Custom | https://doi.org/10.5281/zenodo.19503257 |
| Data synchronization | Custom | https://doi.org/10.5281/zenodo.19503257 |
| Data analysis | Custom | https://doi.org/10.5281/zenodo.19503257 |
| **Other** | | |
| Protocol for Viral Injection | protocols.io | dx.doi.org/10.17504/protocols.io.6qpvrwbd3lmk/v1 |
| Protocol for Lens Implant | protocols.io | dx.doi.org/10.17504/protocols.io.81wgbwj7ygpk/v1 |
| Protocol for EMG Fabrication | protocols.io | dx.doi.org/10.17504/protocols.io.81wgbwj7ygpk/v1 |

| | | |
|---|---|---|
| Protocol for EMG Implant | protocols.io | dx.doi.org/10.17504/protocols.io.e6nvw4w5zlmk/v1 |
| Protocol for Histology | protocols.io | dx.doi.org/10.17504/protocols.io.5qpvodey7g4o/v1 |
| Protocol for Behavior Training | protocols.io | dx.doi.org/10.17504/protocols.io.kqdg31m61l25/v1 |
| Protocol for 2-photon imaging | protocols.io | dx.doi.org/10.17504/protocols.io.4r3l21d2jg1y/v1 |
| Protocol for EMG Recordings | protocols.io | dx.doi.org/10.17504/protocols.io.36wgqpx4yvk5/v1 |
| Protocol for 2-photon Stim. Calibration | protocols.io | dx.doi.org/10.17504/protocols.io.ewov11rqpvr2/v1 |

**Table S6. Key resource table.**
